# Early-life mitochondrial DNA damage results in lifelong deficits in energy production mediated by redox signaling in *Caenorhabditis elegans*

**DOI:** 10.1101/2020.11.25.398164

**Authors:** John P. Rooney, Kathleen A. Hershberger, Elena A. Turner, Lauren J. Donoghue, Laura L. Maurer, Ian T. Ryde, Jina J Kim, Rashmi Joglekar, Jonathan D. Hibshman, Latasha L. Smith, Dhaval P. Bhatt, Olga R. Ilkayeva, Matthew D. Hirschey, Joel N. Meyer

## Abstract

The consequences of damage to the mitochondrial genome (mtDNA) are poorly understood, although mtDNA is more susceptible to damage than nuclear DNA (nucDNA), and many environmental toxicants target the mitochondria. Reports from the toxicological literature suggest that exposure to early-life mitochondrial damage could lead to deleterious consequences later in life (the “Developmental Origins of Health and Disease” paradigm) but reports from other fields often report beneficial (“mitohormetic”) responses to such damage. Here, we test the effects of low (causing no change in lifespan) levels of ultraviolet C (UVC)-induced, irreparable mtDNA damage during early development in *Caenorhabditis elegans*. This exposure led to life-long reductions in mtDNA copy number and steady-state ATP levels, accompanied by increased oxygen consumption and altered metabolite profiles, suggesting inefficient mitochondrial function. Exposed nematodes were also developmentally delayed, reached smaller adult size, and were rendered more susceptible to subsequent exposure to chemical mitotoxicants. Metabolomic and genetic analysis of key signaling and metabolic pathways supported redox signaling during early development as a mechanism for establishing these persistent alterations. Our results highlight the importance of early-life exposures to environmental pollutants, especially in the context of exposure to chemicals that target mitochondria.

## Introduction

The mitochondrial genome (mtDNA) is small (16,569 bases in humans) and encodes a small number of genes: 13 proteins, 22 tRNAs and 2 rRNAs in humans, with similar numbers in most metazoans. Nonetheless, it is critical: the proteins are essential subunits of the mitochondrial respiratory chain (MRC) complexes, while the tRNAs and rRNAs are required for translation of those proteins. The health significance of mtDNA homeostasis is demonstrated by the fact that a large number of diseases are caused by mutations (Chinnery, 2015) or depletion (Suomalainen & Isohanni, 2010) of mtDNA, which is normally present in thousands of copies per cell. In addition to these specific mtDNA-related diseases, mitochondrial dysfunction has been implicated in many more common human diseases including cancer, diabetes, metabolic syndrome and neurodegenerative conditions (Nunnari & Suomalainen, 2012). Further support for the importance of mitochondrial function to health is provided by evidence for toxicity associated with many pharmaceuticals that target mitochondria (Dykens & Will, 2007).

Globally, environmental pollutants are a major environmental driver of loss of years of life, conservatively estimated at three times greater than malaria, AIDS, and tuberculosis combined (Landrigan et al., 2018). Developmental and low-level exposures are of particular concern (Braun & Gray, 2017; Gross & Birnbaum, 2017). The evidence that mitochondria are important targets of many environmental toxicants is growing, yet the consequences of such mitotoxicity are poorly understood (Meyer, Hartman, & Mello, 2018). mtDNA may be a particularly important target, because of the lack of some DNA repair pathways, in particular nucleotide excision repair (NER)(Roubicek & de Souza-Pinto, 2017). Without NER, DNA lesions caused by common environmental stressors such as ultraviolet radiation, polycyclic aromatic hydrocarbons, and mycotoxins are irreparable in mtDNA. *In vitro*, such lesions can impair the progression of the mitochondrial DNA polymerase γ (Graziewicz, Sayer, Jerina, & Copeland, 2004; Kasiviswanathan, Gustafson, Copeland, & Meyer, 2012), potentially resulting in decreased mtDNA copy number or mutagenesis *in vivo*. However, despite the fact that many common environmental exposures cause irreparable mtDNA damage (Meyer et al., 2013), the *in vivo* effects of such damage remain poorly understood.

We have previously employed ultraviolet C radiation (UVC) exposures to generate mtDNA damage that is irreparable due to the absence of NER (nucDNA damage is also caused, but is efficiently repaired). UVC is not environmentally relevant since stratospheric ozone absorbs UVC wavelengths; however, it provides a useful laboratory tool for near-exclusive production of damage (photodimers) that is repaired by NER, compared to UVB and UVA which also produce significant oxidative DNA damage. We reported that early-life (1^st^ larval stage) exposures to such damage reduced levels of mtDNA, mtRNAs, total steady-state ATP levels, and oxygen consumption in developing *C. elegans* (Bess, Crocker, Ryde, & Meyer, 2012; Leung et al., 2013). What might the long-term impact of such mtDNA damage be? Although irreparable, UVC-induced mtDNA damage is slowly removed via a process involving autophagic machinery in both adult *C. elegans* and human cells in culture (Bess et al., 2012; Bess, Ryde, Hinton, & Meyer, 2013). We also found that the nematode mitochondrial DNA polymerase γ, *polg-1*, was dramatically upregulated by UVC exposure (Leung et al., 2013), suggesting the potential for an adaptive response. An adaptive response appears to be consistent with multiple examples of developmental mitochondrial dysfunction resulting in beneficial later-life outcomes in *C. elegans* and other species, including stress resistance and increased lifespan, sometimes termed “mitohormesis” (Dillin et al., 2002; Houtkooper et al., 2013; Mouchiroud et al., 2013; Ng, Ng, van Breugel, Halliwell, & Gruber, 2019; Rea, Ventura, & Johnson, 2007). On the other hand, a growing body of environmental health literature suggests that environmental exposures during development can lead to adverse outcomes later in life, sometimes described as the Developmental Origins of Health and Disease (DOHaD) (Barouki, Gluckman, Grandjean, Hanson, & Heindel, 2012; Grandjean et al., 2015), including a recent example of long-term mitochondrial dysfunction associated with developmental mitochondrial toxicity (Lozoya et al., 2020).

Here, we asked if a low-level, *in vivo*, developmental environmental exposure that causes irreparable mtDNA damage would result in hormetic or adverse later life outcomes related to mitochondrial function, and tested the mechanisms mediating observed changes.

## Results

We employed a previously developed (Bess et al., 2012; Leung et al., 2013) protocol that involves exposing growth arrested L1 larvae to 7.5 J/m^2^ UVC three times (**Fig. 1A**), 24 hours apart, in order to cause a high level of irreparable mtDNA damage while permitting nucDNA repair. Importantly, this is not a highly toxic level of UVC exposure, as demonstrated by the fact that there was no effect on lifespan (**Fig. 1B**). However, this protocol consistently leads to a significant reduction in N2 worm length that is maintained at least 72 hours post-UVC exposure (**Fig. 1C**). We then examined mitochondrial and organismal phenotypes at later timepoints and tested signaling pathways involved in the response to this damage. Many of these studies were conducted using strains (*glp-1* and PE255, an ATP reporter strain created in a *glp-4* background) in which germline proliferation fails at 25° C (Kimble & Crittenden, 2005) in order to exclude the confounding effects of the high level of mtDNA replication that occurs during gametogenesis (Tsang & Lemire, 2002).

**Figure 1.**
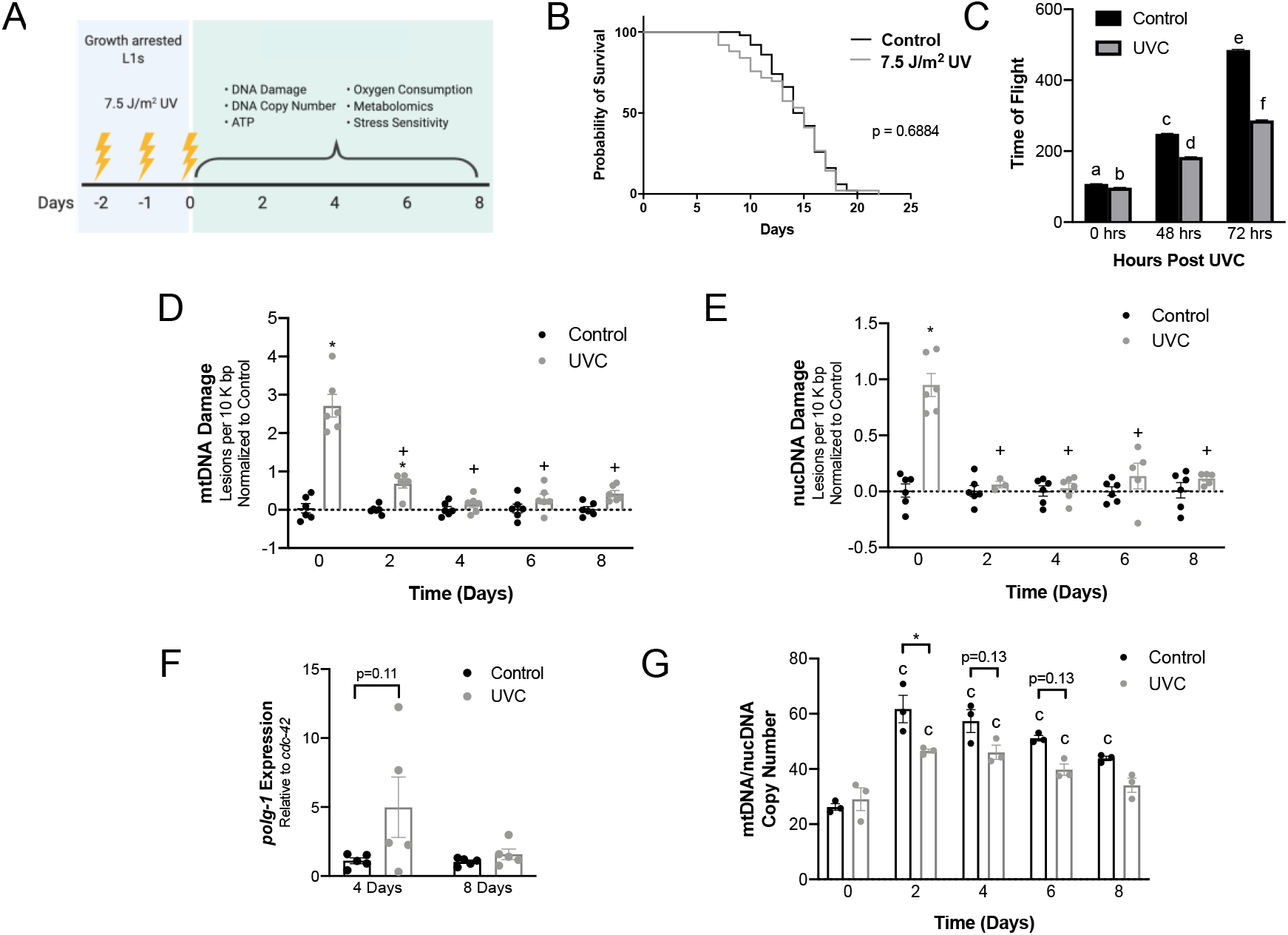
Early-life, irreparable mtDNA damage does not affect lifespan and is removed quickly. **A.** Schematic depicting experimental design. Growth arrested L1s were exposed to UVC radiation three times over 48 hours and transferred to normal culture conditions. Assays were performed at 0-, 2-, 4-, 6-, and 8-days post UVC exposure. Many studies were performed in germline proliferation-deficient strains. **B.** Lifespan is unchanged in N2 nematodes exposed to 7.5 J/m^2^ UVC radiation 3 times over 48 hours as unfed, arrested L1 larvae (p=0.688, Mantel-Cox test). Median lifespan was 15 days for both controls and UVC exposed, while maximal lifespans were 20 days in the controls and 22 days in UVC exposed (20° C). Lifespan days were counted from hatch. n=25-26 per group per experiment. Survival curve combines two independent experiments. **C.** UVC decreases time of flight (TOF; a surrogate for worm length) at 0-, 48-, and 72-hours post-UVC exposure. Two-way ANOVA with Tukey’s correction for multiple comparisons. n > 4000 per group. 2 independent experiments. **D.** UVC induces mtDNA damage that is not repaired but is removed in somatic cells (*glp-1* strain raised at 25° C) by four days post UVC exposure. Data is normalized to control at each timepoint. Two-way ANOVA with Bonferroni’s correction for multiple comparisons (9 comparisons). n = 3 per group per experiment (2 independent experiments). * p ≤ 0.05 compared to control at same timepoint. + p ≤ 0.05 compared to UVC at day 0. **E.** nucDNA damage does not accumulate and is quickly repaired. Two-way ANOVA with Bonferroni’s correction for multiple comparisons (9 comparisons). n = 3 per group per experiment (2 independent experiments). * p ≤ 0.05 compared to control at same timepoint. + p ≤ 0.05 compared to UVC at day 0. **F.** Induction of transcription of the mtDNA polymerase γ (*polg-1*) is large but variable at 4 days post exposure in the UVC exposed animals compared to control animals. Two-way ANOVA with Tukey’s multiple comparisons correction. n=5 per group. **G.** mtDNA/nucDNA copy number ratio in UVC exposed *glp-1* nematodes is significantly lower than control animals at 2 days post-exposure and has a strong trend of remaining reduced for the length of the experiment. Two-way ANOVA with Bonferroni’s correction for multiple comparisons (13 comparisons). n = 1 per group per experiment (3 independent experiments). * p ≤ 0.05. “c” indicates p ≤ 0.05 compared to control at day 0.

### mtDNA damage accumulates over consecutive UVC exposures

We first tested the persistence of UVC-induced mtDNA damage during the worms’ lifespan. UVC-induced DNA damage largely comprises photodimers and 6,4-photoproducts, both of which consist of covalent bonds between adjacent bases; we use the term “photolesion” in this manuscript to describe this class of DNA damage. Although photolesions are not repaired in mtDNA, they are slowly and incompletely removed by autophagy in adult nematodes (Bess et al., 2012); furthermore, damaged genomes may be diluted by the extensive mtDNA replication that occurs during development (Tsang & Lemire, 2002). We measured significantly higher levels of mtDNA damage immediately after the third and final dose of UVC exposure (time 0 days) compared to the non-exposed controls at the same time point (**Fig. 1D** **and Fig. S1A)**. mtDNA damage in UVC exposed animals at day 2 was significantly higher than their non-exposed controls. All other data points showed no distinguishable differences between amount of mtDNA damage in control and UVC-exposed animals at the same time point. This suggests that mtDNA damage is either removed rapidly via mechanisms of mitochondrial quality control, which seems unlikely given the slow kinetics of photodimer removal (Bess et al., 2012), or the mtDNA damage is diluted due to rapid proliferation of mitochondria during development (Leung et al., 2013). The apparent (though not significant) slight increase in mtDNA damage in UVC-exposed animals at day 8 could be a result of oxidative damage resulting from mitochondrial reactive oxygen species (ROS) production, however, this assay cannot distinguish between different types of DNA damage. A significant increase in nucDNA lesions in UVC-exposed compared to control animals was observed at day 0, but not at any other time point. (**Fig.1E** **and Fig. S1B)**. Notably, there was approximately 3-fold higher mtDNA damage in UVC-exposed animals at day 0 compared to nucDNA damage, indicating that nucDNA photolesion damage was removed rapidly between arrested L1 UVC exposures. This replicates our previous observations (Bess et al., 2012; Leung et al., 2013) of much faster photolesion removal in nucDNA, presumably largely from NER.

### Reduction in mtDNA copy number is observed throughout life following developmental UVC exposure

To further characterize DNA integrity following UCV exposure, we analyzed mitochondrial and nuclear DNA copy number. Previous studies have shown that UVC-exposure induces an 8-fold increase in *polg-1* (mitochondrial DNA polymerase γ) expression 48 hours after UVC-exposure (Leung et al., 2013). We confirmed that *polg-1* expression shows a strong trend of remaining elevated up to 4 days post-exposure (p = 0.11) and returns to control levels of expression by 8 days post-exposure (**Fig. 1F**). We hypothesized that photolesions in mtDNA would block the progress of POLG-1 (Kasiviswanathan et al., 2012), resulting in reduced mtDNA content at early timepoints post-exposure. Additionally, because mtDNA damage is not significantly different in control versus UVC-exposed worms in adulthood (**Fig. 1D**), we expected no difference in mtDNA copy number between groups in adulthood. Instead, mtDNA copy number per worm (**Fig. S1C**) and mtDNA/nucDNA ratio **(Fig. 1G)** displayed strong trends of reduction throughout life in the UVC-exposed animals. This was unexpected given the large proliferation of mtDNA copies during development and suggests that there could be other persistent effects of developmental exposure to UVC. There was no effect on nucDNA copy number **(Fig. S1D)**. We observed similar results for the PE255 *glp-4(bn2)* strain **(Fig. S1E)**, used to permit comparisons to ATP levels (below).

### Developmental mtDNA damage did not result in detectable changes in mtRNA transcription in adults

We next asked whether this reduction in mtDNA copy number, and/or the presence of photolesions that might block the mtRNA polymerase (Cline, 2012), would affect the levels of RNAs for mitochondrial proteins, or the stoichiometry of ETC proteins. All ETC complexes except complex II contain proteins coded in both the mitochondrial and nuclear genomes, and proper stoichiometry depends on coordinated production of proteins from both genomes. Improper stoichiometry results in protective mitochondrial unfolded protein responses (UPR^mt^) (Shpilka & Haynes, 2018), which can be triggered by reduced transcription of mtDNA-encoded ETC subunits (Houtkooper et al., 2013). To our surprise, mtDNA-encoded transcript levels in UVC-treated nematodes were unchanged at 4- and 8-days post-exposure **(Fig. S2A)**. This lack of decreased transcription in the UVC-exposed animals might result from the multiplicity of mtDNAs buffering against transcriptional blocks present in a subset of mtDNAs, possibly in combination with compensatory increased transcription. However, a series of experiments designed to detect a compensatory transcriptional response indicated that there were no differences in transcription between control and UVC-exposed animals (**Fig. S2B-C**).

Additionally, we tested for, but did not detect, activation of the UPR^mt^ either transcriptionally **(Fig. S2D**) or at the protein level (data not shown). The *atfs-1* mutant (which is deficient in mounting a UPR^mt^) was protected from larval growth inhibition (**Fig. S2E**), suggesting ATFS-1 mediated transcriptional response is important for inducing growth delay following UVC exposure. Finally, we asked whether exposure to chloramphenicol and doxycycline (two chemicals that inhibit mitochondrial protein translation and nematode larval development (Tsang, Sayles, Grad, Pilgrim, & Lemire, 2001)), would exacerbate UVC-mediated inhibition of larval growth (Bess et al., 2012). We observed that both chloramphenicol and doxycycline further inhibited larval development after UVC exposure (**Fig. S2F**), indicating that, by itself, UVC-induced mtDNA damage does not entirely abrogate translation of proteins coded in mtDNA. Overall, this data suggests that mtDNA damage via UVC exposure, even at this relatively high level of damage, does not appear to result in overall shifts in transcriptional response of the mitochondrial or nuclear genomes, and that mitochondrial protein translation remains functional.

### Mitochondrial respiration was altered in response to developmental mtDNA damage

Given that mtDNA copy number was reduced, but RNAs of mitochondrial proteins were expressed normally after UVC exposure, we next asked if mitochondrial function in later life was impacted by an early life exposure to UVC. To directly assess mitochondrial function, we used the Seahorse XF Analyzer to measure basal, maximal and non-mitochondrial respiration rates at days 4- and 8- post-UVC *in vivo* (A.L. Luz, Smith, Rooney, & Meyer, 2015).

Unexpectedly, basal oxygen consumption rates were significantly increased in UVC-exposed nematodes at 4 days post exposure, but not at 8 days post-exposure **(Fig. 2A)**. Further, basal respiration at 8 days post-exposure were significantly reduced in both the control and UVC exposed groups compared to basal respiration at 4 days post-exposure, consistent with decreased basal respiration with increased age. At 4 days post-exposure, the mitochondrial uncoupler FCCP increased oxygen consumption in non-exposed animals 2.2-fold but had no effect in UVC-exposed nematodes (**Fig 2B)**. Spare capacity (the difference of maximal and basal respiration) was significantly lower in UVC-exposed animals compared to non-exposed animals at 4 days post-exposure (**Fig 2C**), indicating that UVC-exposed animals were operating at or near maximal respiration. Interestingly, non-mitochondrial respiration was similar in the exposed and non-exposed groups at 4 days post-exposure, but there was a significant increase in non-mitochondrial oxygen consumption in the UVC-exposed animals at 8 days post-exposure (**Fig 2D**). Taken together, this data indicates that 4 days after an early life exposure to UVC, basal respiration is increased to near maximal compared to non-exposed controls at the same time point. However, at 8 days post-exposure, there is no difference between exposed and non-exposed groups with respect to basal and maximal respiration, yet there is a significant increase in non-mitochondrial respiration in the exposed compared to non-exposed groups.

**Figure 2.**
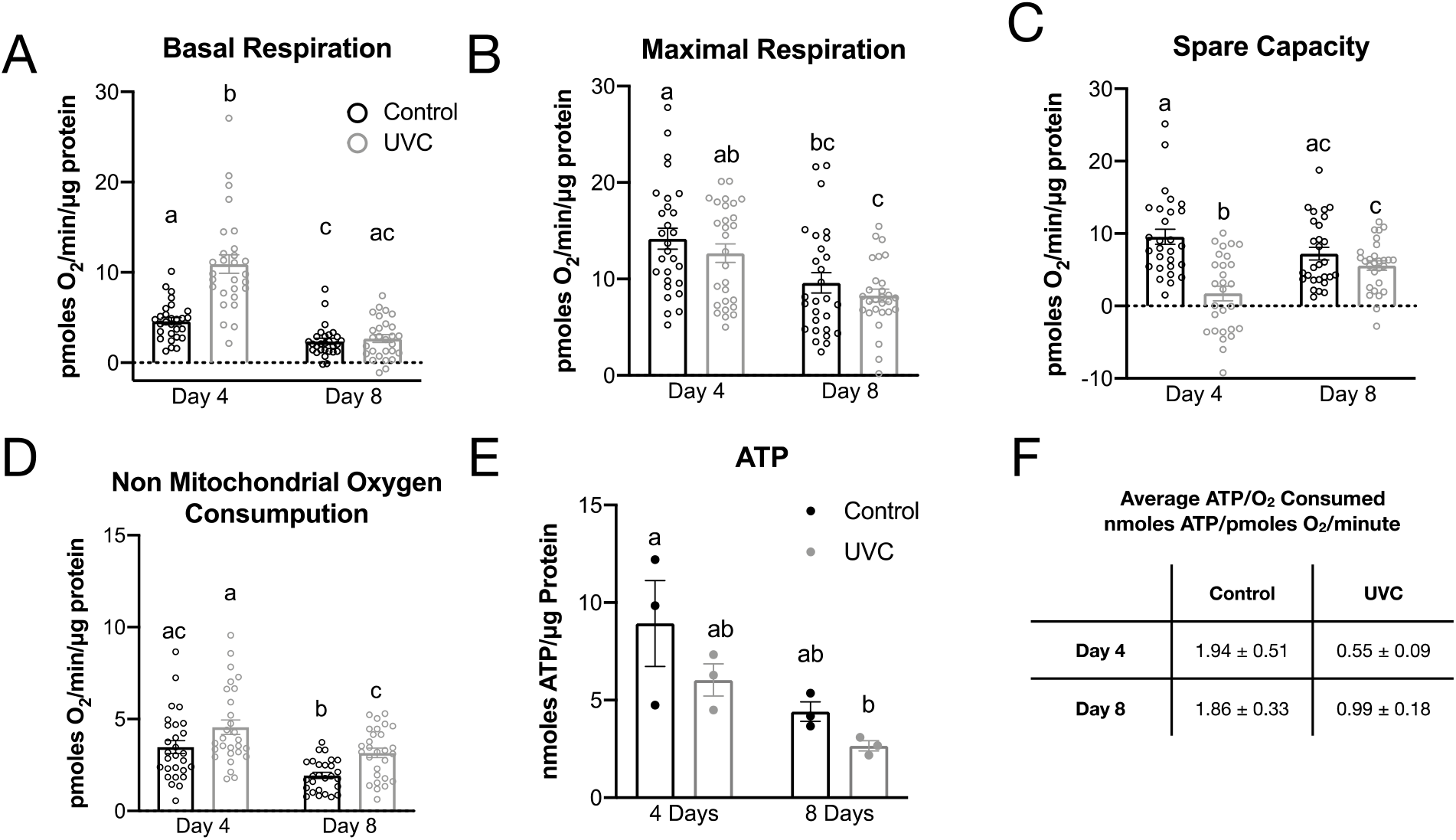
Early-life mtDNA damage results in deficient mitochondrial respiration and decreased ATP per oxygen consumption. **A.** Basal respiratory rates were significantly increased in UVC-exposed nematodes at 4 days post-exposure, but not at 8 days. **B.** Maximal respiration was unchanged between control and UVC treated groups but decreased with age. **C.** Spare capacity was significantly decreased in UVC exposed animals 4 days after exposure and returned to control levels by 8 days post-exposure. **D.** Non mitochondrial oxygen consumption was significantly higher in UVC exposed animals compared to control animals at 8 days post-exposure. For data shown in A-D: Data are presented as mean with error bars representing the standard error. Open circles are individual data points (4 independent experiments, n = 7 per group per experiment). Letters show which groups are significantly different (p ≤ 0.05). Two-way ANOVA with Tukey’s correction for multiple comparisons. **E.** UVC exposure leads to a trend of reduced steady-state ATP in *glp-1*(q244). Two-way ANOVA with Tukey’s correction for multiple comparisons. n=3 per group. **F.** The ratio of ATP per oxygen consumed was calculated by dividing the average steady-state ATP value by the average basal respiration value for each group.

To further analyze the functional consequences of increased basal oxygen consumption rates in the 4-day post-UVC exposed animals, we measured steady-state ATP levels using two methods. ATP levels in extracts from *glp-1* nematodes at 4- and 8- days post-UVC exposure showed a strong trend of reduction compared to their relative controls **(Fig. 2E)**. Consistent with previous results (Braeckman, Houthoofd, De Vreese, & Vanfleteren, 2002), ATP content decreased in an age-dependent fashion by roughly 50% between days 4 and 8 in both control and UVC-exposed animals. We also measured ATP using the *in vivo* ATP reporter strain PE255 and found similar results of decreased steady-state ATP in UVC-exposed animals compared to controls in later life stages (days 6-10 post-exposure) (**Fig. S3A**), with the largest reduction (44%) at day 8. As a complementary measure of energy availability, we measured movement, and found that even lower levels of developmental UVC exposure reduced spontaneous movement in adults 8 days post-exposure (**Fig. S3B**).

The increased basal oxygen consumption and lack of spare respiratory capacity combined with reduced ATP levels in UVC-exposed nematodes suggested persistently inefficient mitochondrial function. The decrease in spare respiratory capacity might be partially explained by the decrease in mitochondrial content suggested by decreased mtDNA content, or by decreased availability of substrate. We also note that although at 8 days post exposure basal oxygen consumption rates were no longer elevated in UVC treated worms, ATP levels still trended lower, suggesting that mitochondrial efficiency was also reduced, resulting in a reduction in ATP levels per unit oxygen consumed (**Fig. 2F**).

### Metabolic flexibility is critical to tolerance of early-life UVC exposure

We hypothesized that in response to inefficient ETC function, UVC-exposed nematodes would shift metabolism towards alternate metabolic pathways. We tested whether strains carrying mutations in two major regulators of energy metabolism (*aak-2,* activated upon increase in the AMP:ATP ratio (Apfeld, O’Connor, McDonagh, DiStefano, & Curtis, 2004) and *nhr-49*, a nuclear hormone receptor that regulates energy metabolism in response to mitochondrial impairment (Van Gilst, Hadjivassiliou, Jolly, & Yamamoto, 2005)) would be sensitized to UVC-induced inhibition of larval development (Bess et al., 2012). *aak-*2 (**Fig. 3A**) and *nhr-49* (**Fig. 3B**) mutant strains were both approximately two-fold more sensitive to early-life UVC exposure, supporting the importance of the nematodes’ capacity to regulate energy metabolism in response to early-life mtDNA damage. Importantly, for extrapolating the results from mutant versus wild-type N2 studies to the outcomes observed in *glp-1* animals used in other experiments in this manuscript, the growth decrease observed in the N2 strain was also observed (to a lesser extent) in the *glp-1* strain (**Fig. 3C**).

**Figure 3.**
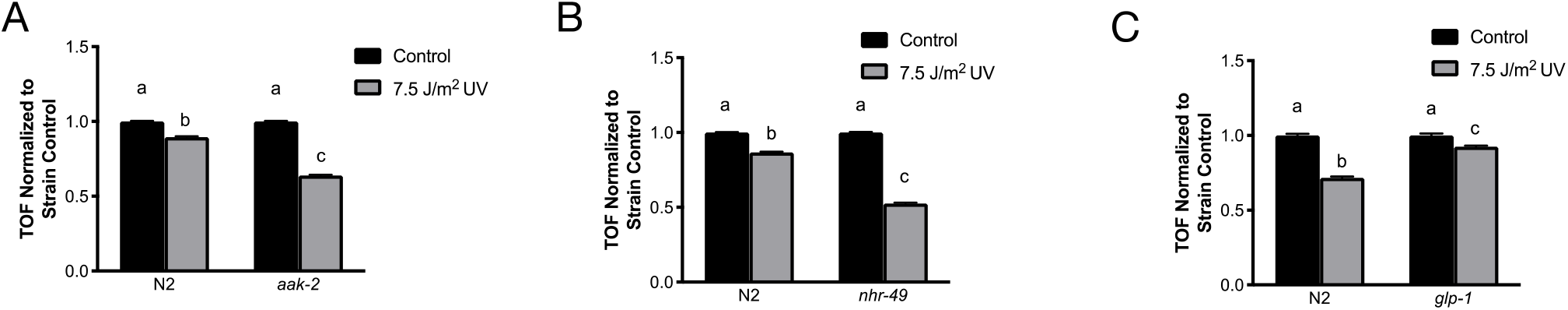
*aak-2* and *nrh-49* mutants are sensitive to UVC. **A.** *aak-2* (36% decrease) and **B.** *nhr-49* (48% decrease) are significantly growth delayed 72 hours post UVC exposure compared to N2 animals exposed to UVC (10-13% decrease). **C.** *glp-1* animals exposed to UVC had a mild growth delay (7.5% decrease) that was less than N2 animals exposed to UVC (30% decrease) in the same experiment. n > 500 in each group from 2 independent experiments. TOF values from UVC exposures are normalized to strain controls. Letters show which groups are significantly different (p ≤ 0.05). Two-way ANOVA with Tukey’s correction for multiple comparisons.

Given this genetic evidence for protective metabolic restructuring, we next tested for alterations in transcript levels of key intermediary metabolic enzymes that regulate TCA cycle metabolism (*pdp-1, pdhk-2*), glycolysis (*gpd-3*), the glyoxylate cycle (*gei-7*), fatty acid B-oxidation (*acs-2*), and gluconeogenesis (*pck-1*), but detected no changes at either 4- or 8-days post-exposure **(Fig. S4)**. However, the lack of altered transcriptional regulation of intermediary metabolic pathways does not rule out significant metabolic alterations, as there are many non-transcriptional mechanisms of enzyme activity regulation. To directly test for metabolic shifts, we next measured levels of organic acids, amino acids and acyl carnitines.

### Targeted metabolomics supports NADPH stress and altered metabolic function in adults

We observed age-related decreases in the levels of most organic acids between days 4 and 8 in both control and UVC-exposed animals (**Fig 4A**). In contrast, fumarate and malate appeared to be elevated in the UVC-exposed animals compared to non-exposed controls and statistical analysis revealed that fumarate was significantly elevated in the UVC-exposed animals compared to controls at 4 days post-exposure (**Fig 4B**). Additionally, there was a significant main effect of increased malate in the UVC-exposed animals compared to controls though multiple comparisons testing did not reveal statistical significance at a specific time point (**Fig 4C**). This increase suggests cataplerosis from the TCA cycle, and activation of the malate-aspartate shuttle, potentially in response to NADPH stress. The NADP^+^/NADPH ratio in the mitochondria can be increased (referred to as “NADPH stress”) through reduction of oxidized glutathione. The malate-aspartate shuttle is one of the principal mechanisms for transferring reducing equivalents into the mitochondria, which can be utilized to decrease this ratio. Amino acid levels also generally declined with age, consistent with reduced overall metabolism in aging nematodes (**Fig. 4D**). Notably, Asx (aspartate/asparagine), was significantly increased in UVC-exposed animals compared to control animals at day 8 post-exposure **(Fig. 4E**). This observation also supports the hypothesis of activation of the malate-aspartate shuttle. Lastly, a number of long chain acyl-carnitines (LCAC) were elevated in UVC samples at day 4 **(Fig. S5),** which is often indicative of defects in, or altered regulation of, fatty acid beta-oxidation. When absolute LCAC levels were compared UVC-related increases were seen in both C18 and C18:1 species, which are relatively abundant (data not shown).

**Figure 4.**
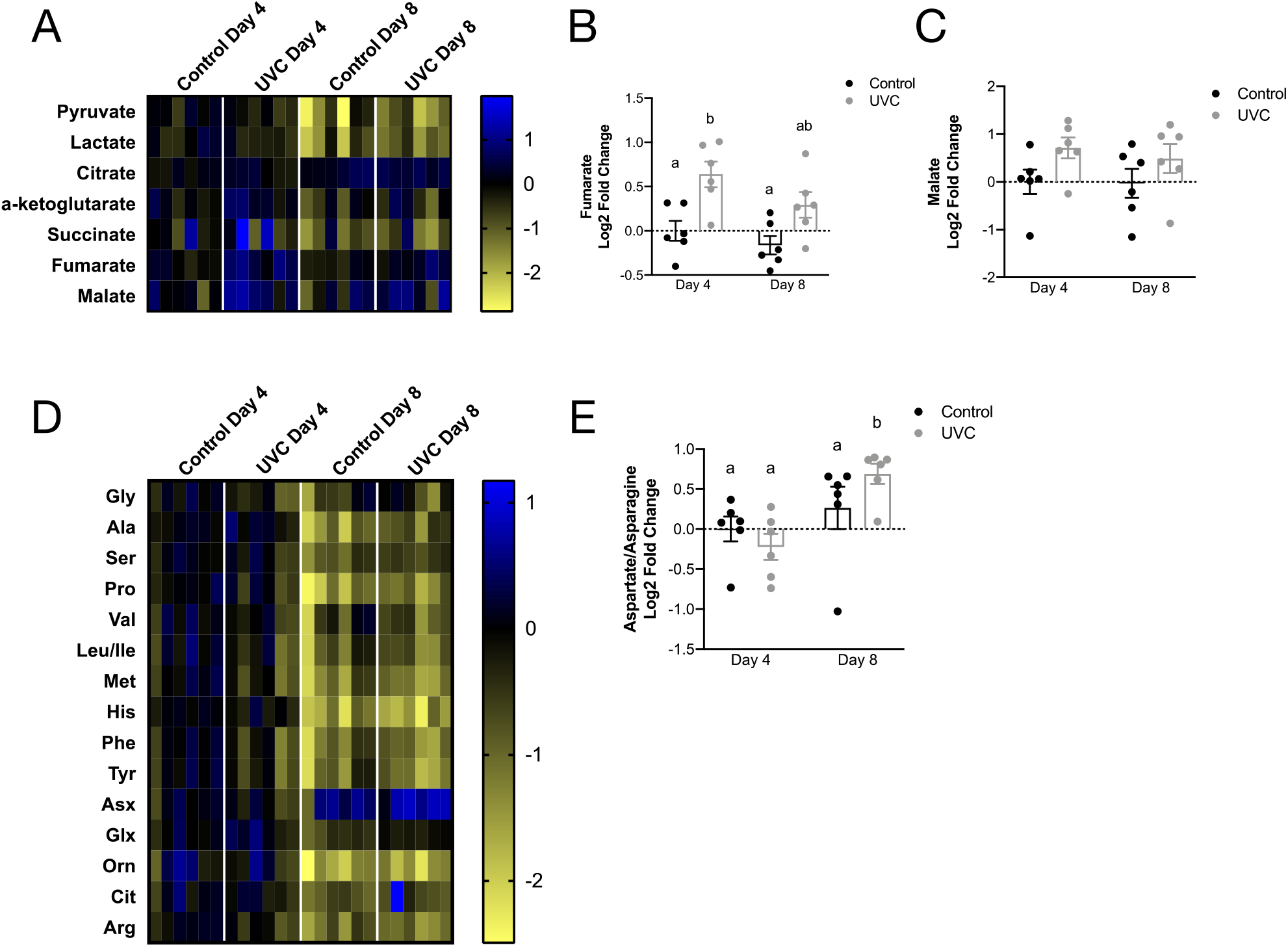
Metabolomics analysis suggests that the malate-aspartate shuttle is activated after UVC exposure. **A.** Heat map of organic acids at 4- and 8-days post-exposure in control and UVC exposed animals. **B.** Fumarate is significantly increased in UVC exposed animals at 4 days post-exposure. **C.** Malate trends toward increased in UVC exposed compared to control animals at 4- and 8-days post-exposure. **D.** Heat map of amino acids at 4- and 8-days post-exposure in control and UVC animals. Generally, amino acid levels decline with age with no effect of UVC exposure. **E.** Asx (aspartic acid/asparagine) is significantly increased in UVC-exposed animals compared to controls at 8-days post-exposure. Error bars represent standard error. Letters show which groups are significantly different (p ≤ 0.05). Two-way ANOVA with Tukey’s correction for multiple comparisons. n = 6 per group, 1 independent experiment.

### Mitochondrial SOD mutants are more sensitive to UVC exposure

The data shown thus far provides compelling evidence for NADPH stress as a mechanism of later life effects of early-life exposure to UVC. Specifically, we observe increased basal oxygen consumption (in the context of reduced ATP and movement) and evidence of increased flux through the malate-aspartate shuttle. Mitochondrial dysfunction, particularly in the context of increased overall oxygen consumption, can result in increased oxidative protein damage. We observed a significant increase in carbonyl groups of cytoplasmic (8%) and mitochondrial proteins (20%) 4 days post-UVC exposure **(Fig. 5A-B)**, suggesting increased ROS production later in life in response to early-life mtDNA damage.

**Figure 5.**
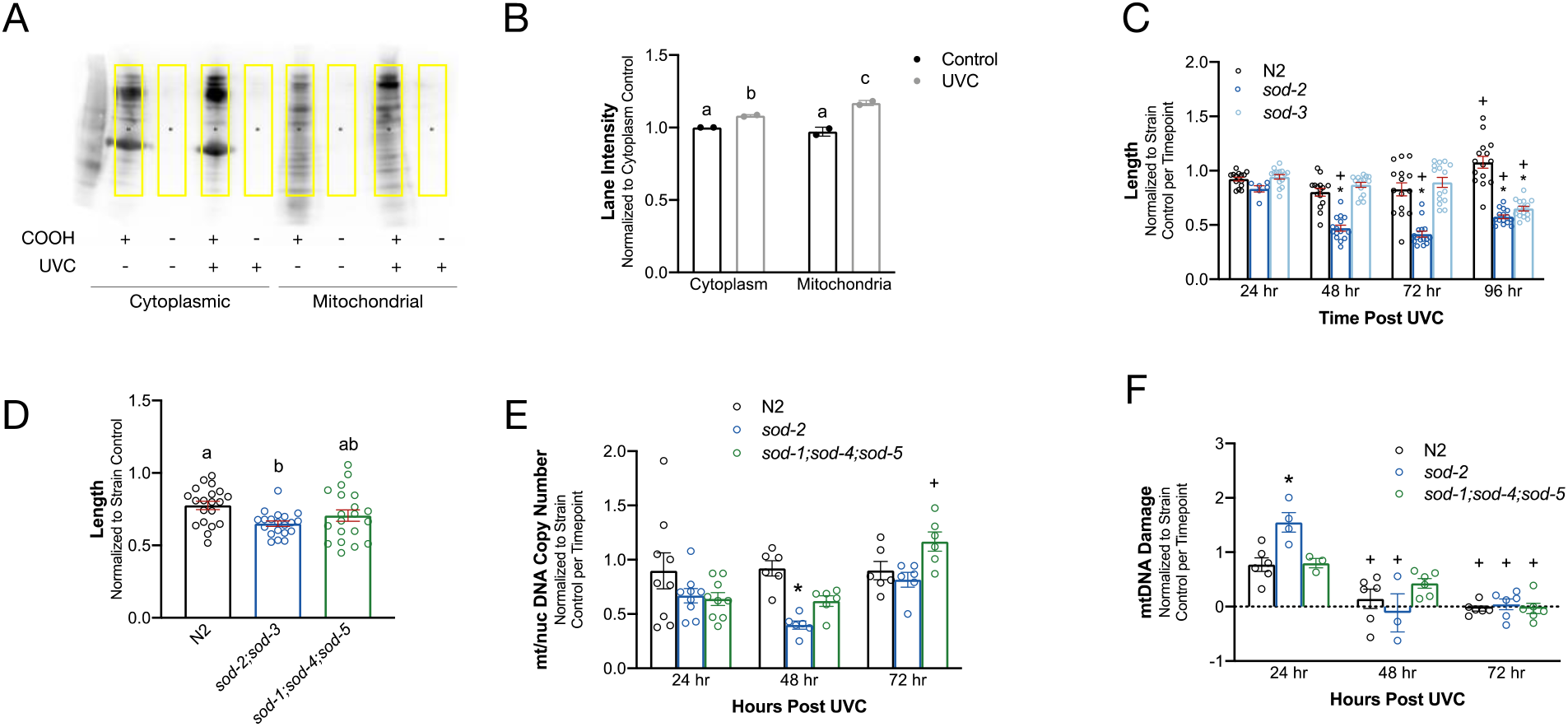
Mitochondrial SOD mutants are more sensitive to UVC exposure. **A.** Representative image of oxyblot at 4 days post-exposure demonstrates increased oxidative protein damage in UVC exposed nematodes in both cytoplasmic and mitochondrial fractions. Yellow boxes indicate area of the blot that was quantified. COOH = derivatization reagent (required for detection), UVC = UVC exposure. **B.** Quantification of oxyblot. Error bars represent standard error. Letters show which groups are significantly different (p ≤ 0.05). Two-way ANOVA with Tukey’s correction for multiple comparisons. n = 2 per group, 2 independent experiments. **C.** *sod-2* and *sod-3* mutants are more sensitive to UVC induced growth delay 96 hours post-exposure. Each UVC exposed strain is normalized to its own non-exposed control at each time point. Two-way ANOVA with Bonferroni correction for multiple comparisons (17 comparisons). n = 6–15 per group. * p ≤ 0.05 compared to N2 UVC exposed at same time point. + p ≤ 0.05 compared to 24 hr UCV exposed of same strain. **D.** The mitochondrial SOD double mutant *sod-2;sod-3*, is more sensitive to UVC induced growth delay 72 hours post UVC exposure than wild type N2 though the cytosolic and extracellular combination knockout (*sod-1;sod-4;sod-5*) is not. One-way ANOVA with Tukey’s correction for multiple comparisons. n = 20 per group. Different letters indicate a significant difference (p ≤ 0.05) between groups **E**. mtDNA:nucDNA ratio in SOD mutants over time. Each UVC exposed strain is normalized to its own non-exposed control at each time point. Two-way ANOVA with Bonferroni correction for multiple comparisons (12 comparisons). n = 6–9 per group. * p ≤ 0.05 compared to N2 UVC exposed at same time point. + p ≤ 0.05 compared to 24 hr UCV exposed of same strain. **F**. mtDNA damage in SOD mutants. Each UVC exposed strain is normalized to its own non-exposed control at each time point. mtDNA damage was increased by more than two-fold in UVC treated *sod-2* nematodes in comparison to both UVC-treated wild-type and *sod-1;sod-4;sod-5* mutant nematodes at the 24 hour timepoint. By 48 h and 72 h, damage levels had returned to our limit of detection in all strains. UVC exposed *sod-1;sod-4;sod-5* mutants were not statistically distinguishable from UVC exposed N2s in terms of mtDNA damage at any timepoint. Two-way ANOVA with Bonferroni correction for multiple comparisons (12 comparisons). n = 3–6 per group. * p ≤ 0.05 compared to N2 UVC exposed at same time point. + p ≤ 0.05 compared to 24 hr UCV exposed of same strain.

The ETC is a major source of intracellular ROS, and mitochondrial dysfunction can result in increased ROS production (M. P. Murphy, 2009), including both superoxide anion (O_2_^−^) and hydrogen peroxide (H_2_O_2_). Mitochondrial ROS may cause damage or serve as mitochondrial signaling molecules (Blajszczak & Bonini, 2017). To investigate the potential functional importance of increased mitochondrial ROS generation, we tested the sensitivity to developmental UVC exposure of nematode strains carrying loss-of-function mutations in all 5 superoxide dismutase (SOD) enzymes (*sod-1, sod-2, sod-3, sod-4,* and *sod-5*), which convert O_2_^−^ to H_2_O_2_ (McCord & Fridovich, 1969). We additionally looked at the effects of UVC exposure on the double *sod-2;sod-3* mutant (mutations in both mitochondrial SOD proteins), and the triple *sod-1;sod-4;sod-5* mutant (mutations in all cytosolic and extracellular SOD proteins).

UVC-induced larval delay was exacerbated in both mitochondrial SOD mutants (*sod-2* and *sod-3*), with the delay occurring as soon as 48-hours post-exposure in *sod-2* nematodes and by 96-hours post-exposure in *sod-3* nematodes **(Fig. 5C)**. Furthermore, the double *sod-2;sod-3* mutant was significantly more growth delayed than wild-type 72 hours after UVC exposure (p = 0.011), but the triple *sod-1;sod-4;sod-5* mutant was not (p = 0.229) **(Fig. 5D)**. Interestingly, the *sod-2* mutant was more sensitive than the *sod-2;sod-3* mutant, consistent with a previous report (Gruber et al., 2011). Based on this data, we used the *sod-2* and *sod-1;sod-4;sod-5* mutants to examine the role of mitochondrial and cytosolic superoxide dismutase proteins in the response to UVC exposure. We confirmed the growth delay at the molecular level by measuring mitochondrial and nuclear DNA content in wild-type and SOD mutants after UVC exposure.

There was a significant delay in mtDNA:nucDNA copy number expansion in the *sod-2* worms, but not the *sod-1;sod-4;sod-5* worms at 48-hours post-exposure (**Fig. 5E)**. The reduction in this ratio was due to the reduction in mtDNA copy number expansion (**Fig. S6A**) as nucDNA copy number expansion was comparable in all three nematode strains post-UVC exposure (**Fig. S6B)**. mtDNA damage at 24-hours post-exposure was significantly increased in the *sod-2* mutants compared to N2, but not in *sod-1;sod-4;sod-5* mutants (**Fig. 5F**). We detected no statistically significant differences between strains in nucDNA damage (data not shown). In response to UVC, the *sod-2* mutants show increased mtDNA damage 24-hours post-exposure and exhibit a more severe growth delay compared to N2 animals, suggesting a role of mitochondrial ROS production in response to UVC damage.

Given the evidence of increased mitochondrial ROS, we looked at transcript levels of heat shock proteins in the *N2*, *sod-2*, and *sod-1;sod-4;sod-5* strains to test for evidence of the heat shock response contributing to the increased mitochondrial ROS (Agarwal & Ganesh, 2020). However, we did not detect any *sod-2*-specific increases in any tested mitochondrial (*hsp-6, hsp-60a,* and *hsp-60b*), cytosolic (*hsp-16.12* and *hsp-16.41*), or endoplasmic reticular (*hsp-4*) heat shock protein (**Fig. S6C–H**), further supporting NADPH stress as a key player in the increased mitochondrial ROS in response to UVC exposure.

### *Skn-1* mediates response to UVC exposure

The delayed development and indications of increased mtDNA damage following UVC exposure in the mitochondrial, but not cytosolic SOD mutants could be explained in two ways: either mitochondrial O_2_^−^ causes the growth delay by causing oxidative damage; or conversion of mitochondrial O_2_^−^ to H_2_O_2_ is important in reducing the extent of the growth delay via adaptive signaling. Both H_2_O_2_ and O_2_^−^ can cause macromolecular damage, while only H_2_O_2_ can exit the mitochondria to act as a signaling molecule. To address the first hypothesis, we tested whether antioxidant treatment of 10 mM N-acetyl cysteine, 3 mM Trolox, or 1-5 μM MitoQ (all previously shown to protect against various stressors in *C. elegans* (Benedetti et al., 2008; A. L. Luz et al., 2017; W. Yang & Hekimi, 2010; X. Yang et al., 2012) rescued UVC-induced growth delays. We observed no effect of antioxidant treatment on the UVC-induced growth delay (data not shown). To test the hypothesis that redox signaling is protective after UVC exposure, we examined the sensitivity of nematodes deficient in *skn-1,* the nematode homologue of the mammalian oxidant-responsive Nrf2 transcription factor. Nrf2 controls the expression of genes related to detoxification, antioxidant defenses, and intermediary metabolism (Hayes & Dinkova-Kostova, 2014), and nematodes lacking *skn-1* are oxidative stress-sensitive (J. H. An & Blackwell, 2003). Remarkably, *skn-1* mutant nematodes showed only a 7% decrease in growth with UVC exposure **(Fig. 6A)**, suggesting that the dramatic growth delay observed in wild-type nematodes after UVC exposure is partially dependent on the *skn-1* mediated signaling response to ROS, not oxidative stress. We investigated the sensitivity of other transcription factors that mediate the ROS stress response. Nematodes with deletions in *daf-16* (the nematode FOXO transcription factor (Lin, Dorman, Rodan, & Kenyon, 1997; C. T. Murphy et al., 2003)) and *ced-4* (the nematode homologue of Apaf1 (Yee, Yang, & Hekimi, 2014)) showed a similar or more dramatic growth delay compared to the N2 nematodes, indicating that the response of these two transcription factors to UVC exposure does not contribute to the growth delay (**Fig. S7A and B**). Additionally, nematodes with genetically compromised ETC function and increased oxidative stress sensitivity (Dingley et al., 2010) were not more resistant to UVC exposure: both *gas-1* and *mev-1* mutants were more growth delayed than wild type nematodes after UVC exposure **(Fig. S7C and D)**. Together, these data suggest that *skn-1* signaling, and not oxidative damage, is responsible for mediating the response to UVC exposure. Lastly, we tested ATP levels in *skn-1* mutants to determine if *skn-1* signaling was responsible for the persistent phenotypes observed and found that the *skn-1* mutants were no more sensitive to reduction in ATP levels after UVC exposure than N2 **(Fig. 6B),** which is also inconsistent with an oxidative damage theory, since *skn-1* mutants are oxidative stress-sensitive.

**Figure 6.**
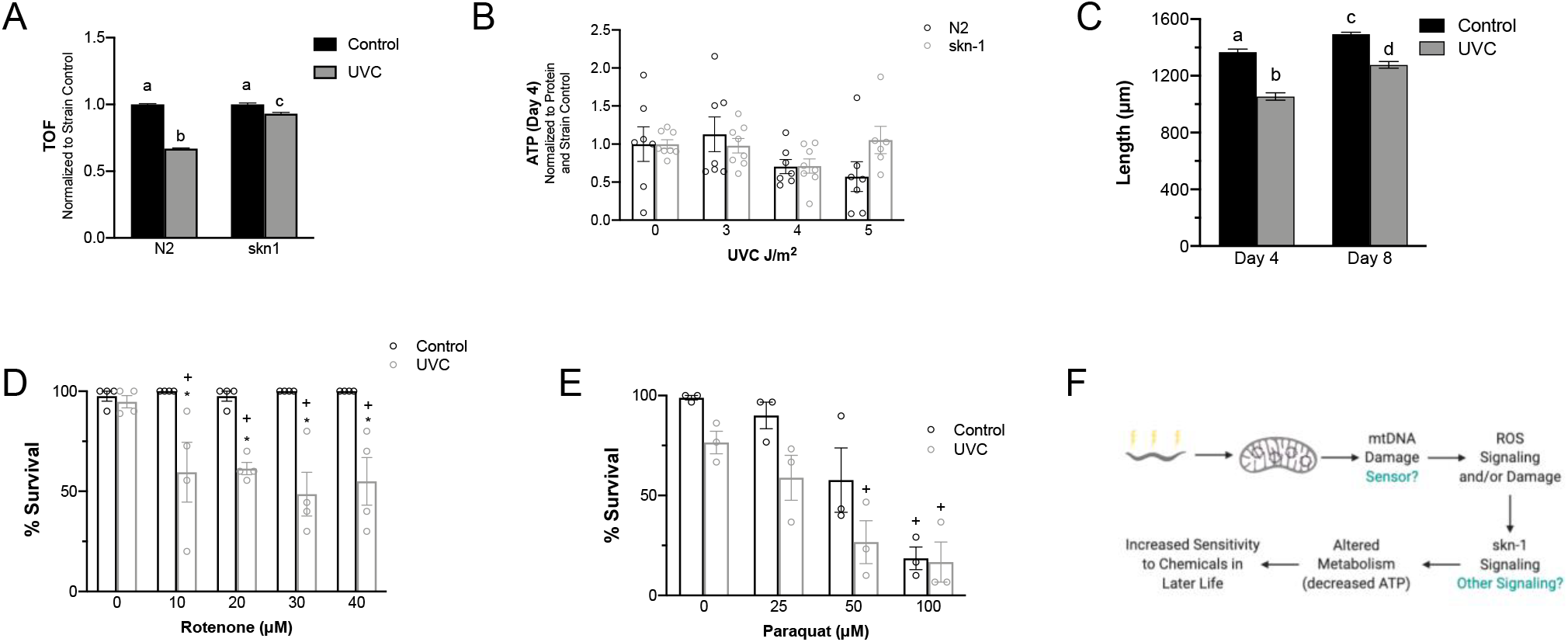
Developmental UVC exposure results in *skn-1* signaling that mediates phenotypes in the adult nematode. **A.** *skn-1* mutants are protected from UVC-induced growth delay 72 hours post UVC exposure. n > 500 in each group from 2 independent experiments. TOF values from UVC exposures are normalized to strain controls. Letters show which groups are significantly different (p ≤ 0.05). Two-way ANOVA with Tukey’s correction for multiple comparisons. **B.** *skn-1* mutants are protected from UVC-induced decrease in ATP levels 4 days post UVC exposure. Two-way ANOVA with Bonferroni correction for multiple comparisons (10 comparisons). **C.** Adult size was reduced in response to UVC at both day 4 and day 8. Letters show which groups are significantly different (p ≤ 0.05). Two-way ANOVA with Tukey’s correction for multiple comparisons. n=60–70 individuals in 2 experimental replicates. **D.** UVC exposed nematodes are more sensitive to the complex I inhibitor rotenone, but not to **E.** paraquat. n=4 per group. Two-way ANOVA with Bonferroni correction for multiple comparisons (13 in D and 10 in E). + p ≤ 0.05 compared to no drug treatment, no UVC exposure control. **F.** Model for ROS involvement in the development of lifelong effects as a result of early-life exposure to UVC. UVC exposure in early life leads to mtDNA damage that is sensed by the cell. This unidentified sensor leads to ROS generated from the ETC. *skn-1* is activated by increased ROS generation, leading to decreased ATP and persistent phenotypes in adults.

### Early life mtDNA damage results in decreased stress resistance in adulthood

As described earlier, there is evidence for both negative and positive effects of mitochondrial dysfunction and toxicity during development. Developmental mtDNA damage resulted in many of the same phenotypes as mitohormesis, suggesting the potential for a common mechanism. However, we found that although adult size was significantly (10–20%) reduced in response to developmental mtDNA damage **(Fig. 6C)** and ATP levels were reduced (**Figs. 2 and S6**), consistent with mitohormetic phenotypes, lifespan was statistically unchanged from that of control nematodes **(Fig. 1B)**, whereas mitohormesis is associated with lifespan extension. Median lifespans were 15 days, and maximal lifespans were 20 and 22 days, in control and UVC treated animals, respectively. However, organisms outside of the laboratory are exposed to a variety of stressors, and there is evidence for “mitohormetic” responses resulting in increased stress resistance (Yun & Finkel, 2014). Instead of increased stress resistance, however, we found that UVC treated nematodes were more sensitive to rotenone **(Fig. 6D)** than their untreated counterparts, though we detected no difference in sensitivity to paraquat (**Fig. 6E**).

## Discussion

### DOHaD or lifespan extension and stress resistance

Our results indicate that irreparable mtDNA damage incurred early in *C. elegans* development can lead to a lifelong reduction of mtDNA:nucDNA copy number, increased basal oxygen consumption and lack of spare respiratory capacity in the context of reduced ATP levels, and sensitivity to later-life chemical challenge, thus supporting the DOHaD hypothesis. These results are in contrast to many others in the literature that find hormetic responses, including increased lifespan and stress resistance, in response to knockdown of ETC genes (Dillin et al., 2002; Durieux, Wolff, & Dillin, 2011; Houtkooper et al., 2013; Rea et al., 2007; Zubovych, Straud, & Roth, 2010) and some chemical exposures (Mouchiroud et al., 2013; Zubovych et al., 2010). While there are several phenotypes shared between reported hormetic responses in the literature and the phenotypes we observed, including small adult size and reduced ATP levels, the lack of hormesis we observed is not entirely unexpected, as there were also differences in observed phenotypes. In particular, activation of the mitochondrial unfolded protein response, which triggers compensatory mechanisms that extend lifespan (Houtkooper et al., 2013), was not observed in our experiments. We also did not observe a change in lifespan. We note that the majority of the treatments reported in the literature to trigger mitohormesis, particularly RNAi, result in major losses of protein or function. In our experiments, it appears that transcripts and proteins coded in the mtDNA were produced at sufficient levels despite the presence of transcription-blocking photolesions, presumably due to the multiplicity of mtDNAs. It may be that most environmental chemical exposures belong to a mechanistic category distinct from and possibly poorly modeled by gene knockdown, gene knockout, or specific pharmacological mitochondrial poisoning, because most environmental mitotoxicants are relatively non-specific (Meyer et al., 2018).

More broadly, we suggest that observations of “mitohormesis” should be interpreted cautiously. RNAi experiments are different than what is seen in most human mitochondrial diseases, most of which result from mutations in mitochondrial gene, rather than reductions in transcript and protein levels. That mutations and RNAi knockdown are not identical is further underscored by the fact that in *C. elegans*, many strains that harbor mutations in genes that when knocked down with RNAi result in lifespan extension, actually have reduced lifespans (Rea et al., 2007). Nonetheless, some mutations in ETC subunit genes do lead to lifespan extension. For example, an *nduf-7* (complex 1) mutation increases ROS, activates the UPR^mt^ and extends lifespan (Rauthan & Pilon, 2015). Another report indicates that relatively smaller perturbations of mitochondrial function can lead to later-life, mitohormetic effects by depressing the normally programmed repression of the heat shock response in adulthood (Labbadia et al., 2017). Possible explanations for our differing results include: 1) different mechanisms of mitochondrial toxicity or different mutation-induced forms of protein dysfunction may have different long-term consequences; 2) examination of different health-related endpoints may reveal apparently hormetic vs deleterious consequences; 3) exposure level may be critical: i.e., it is possible that even lower levels of mtDNA damage than those we employed would cause a more classically hormetic response. Both toxicological (Calabrese, 2013) and gene knock-down (Rea et al., 2007) models of hormesis propose beneficial effects only at low levels of inhibition or dysfunction. Another caveat with interpreting model organism hormesis results is suggested by the fact that lifespan extension in *C. elegans* is not always directly relevant to humans, as mutations in *C. elegans* mitochondrial genes that increase lifespan can lead to shortened lifespan in humans (Rea et al., 2007). Finally, there are tradeoffs associated with extending lifespan, bringing into question the apparently unambiguously positive connotation of the term “mitohormesis.” Reduced fertility, smaller adult size, and reduced movement are often seen in conjunction with lifespan extension. These results are consistent with data from recent toxicological studies which have provided evidence for later-life, adverse effects of mitochondrial toxicity. Perhaps most prominently among such reports, developmental exposure to nucleoside reverse transcriptase inhibitors, which interfere with mtDNA homeostasis, results in later-life mtDNA mutations, reduced mtDNA copy number, and reduced mitochondrial function in humans (Poirier, Gibbons, Rugeles, Andre-Schmutz, & Blanche, 2015), rodents (Liu, Nguyen, Baris, & Poirier, 2012) and primates (Liu et al., 2016). Other examples include latent mitochondrial myocardial toxicity seen in patients receiving doxorubicin as chemotherapy (Berthiaume & Wallace, 2007), latent liver carcinogenicity in mice after short term exposure to dichloroacetate (Wood et al., 2015), and increased rates of mitochondrial diseases later in life among survivors of childhood cancers (Hudson et al., 2013).

### A role for mitochondrial reactive oxygen species in signaling and metabolic programming

One of the most striking effects observed in response to early life mtDNA damage was the persistent reduction in steady state ATP levels, especially since this persisted even after mtDNA lesions were removed, and mtDNA-encoded ETC transcript levels were at or above those in untreated nematodes. mtDNA copy number was reduced at this time, but only approximately 20%, which by itself is not sufficient to reduce ATP levels in *C. elegans* (A. L. Luz & Meyer, 2016). Similarly, the lack of change in mtRNA levels or activation of UPR^mt^, suggest that the changes in ATP levels and other phenotypic effects result from signaling, rather than being direct effects of mtDNA damage or copy number changes.

Increased protein oxidation, increased oxygen consumption despite decreased ATP levels and movement, and the reduced sensitivity of *skn-1* mutants to developmental mtDNA damage all support mtROS signaling as a mechanism for the regulation of growth delay **(Fig. 6F)**. Increased mtROS is also consistent with metabolomic shifts suggesting activation of the malate-aspartate shuttle, which can be utilized to move malate into or out of the mitochondria. In the cytosol, malic enzyme can convert malate to pyruvate, generating NADPH from NADP^+^ in the process. However, the malate-aspartate shuttle can also move NADH into the mitochondria, where reducing equivalents can be transferred from NADH to NADP^+^ via the nicotinamide nucleoside transhydrogenase (NNT) using energy from membrane potential. Mitochondrial NADPH is required to reduce glutathione, a key antioxidant that exists as a separate pool from cytoplasmic glutathione (Meredith & Reed, 1982), such that its reduction largely occurs within the mitochondria. In the context of oxidative stress resulting from UVC-induced mtDNA damage, much of the reducing equivalent pool in the mitochondria could be used for reducing oxidized glutathione, making less available for OXPHOS and ATP production, while at the same time reducing the mitochondrial membrane potential.

Our data suggest that developmental UVC exposure results in a low level of mtROS that plays two roles, both causing damage and acting as signaling molecules, with a dominant long-term role for alterations in developmental ROS signaling. ROS play an important signaling role during development, and disruption of this signaling can alter differentiation and proliferation (Hernandez-Garcia, Wood, Castro-Obregon, & Covarrubias, 2010). Short term exposure to H_2_O_2_ in young nematodes results in many similar outcomes to those seen here, including reduced movement and ATP levels, though these effects are transient (Kumsta, Thamsen, & Jakob, 2011). We hypothesize that in our experiments, aberrant ROS generation during development led to *skn-1*-mediated metabolic alterations that program the developing nematode for life. In wild-type worms, the toxic effects of O_2_^−^ are ameliorated by the action of SODs, and thus insufficient to cause developmental delay, as indicated both by the sensitivity of the mitochondrial SOD mutants and by the inability of pharmacological antioxidants to rescue growth delay.

Interestingly, the UPR^mt^, while not induced in wildtype nematodes, was activated in a delayed fashion in *sod-2* mutants, further indicating that mtROS were insufficient in the wild-type background to activate a damage-associated developmental delay. mtROS, however, appears to be required for a SKN-1-mediated growth delay and metabolic remodeling, as indicated by the initially counterintuitive finding that *skn-1* mutants are protected from developmental delay, and by the absence of a reduction in ATP levels in UVC-exposed *skn-1* mutants. More broadly, given that ROS are both damaging and signaling molecules, it is not surprising that alterations in either direction of their tightly regulated developmental levels may have short-term (i.e., growth) and long-term (i.e., metabolic programming) consequences.

### Unexpected mtDNA damage and copy number dynamics

The patterns of change in mtDNA copy number and mtDNA damage were also unexpected in a number of ways. With regards to mtDNA damage, the intriguing reappearance of mtDNA lesions 8 days post-exposure suggests that mtDNA damage may accumulate more quickly with aging in the context of early life exposure than in control animals (in which we did not detect damage at 8 days), possibly due to persistently increased oxidative stress or reduced repair capacity due to inhibited ATP or nucleotide availability.

The complete (at least within our limits of detection) removal of photolesions from mtDNA in developing larvae is in contrast to previous work from our lab in somatic cells of adult nematodes, which had shown just under 40% damage removal after 72 hours in adult nematodes (Bess et al., 2012). mtDNA replication appears to stop or at least fall below detection in adult nematodes, and copy number slowly declines with a half-life of approximately 10 days (Rooney et al., 2014). This half-life roughly corresponds to the rate at which damage is removed, suggesting that mtDNA damage removal in adult nematodes is tied to turnover of mitochondrial genomes. In contrast, mtDNA replication is essential and extensive during development. We hypothesize that increased mtDNA replication (leading to dilution of damaged genomes with new, undamaged copies) and, potentially, more frequent turnover leads to more complete reduction in per-nucleotide levels of damage during development. Supporting this hypothesis, the increase in mtDNA copy number between days 0 and 2 (3.88-fold) is proportionally similar to the reduction in lesion frequency over the same time period (3.98-fold). This would suggest that photodimers in mtDNA are not removed in any quantitatively important fashion during the first 2 days of *C. elegans* development.

With regard to mtDNA copy number, complete lesion removal and induction of *polg-1* suggest that mtDNA replication should be unimpeded, yet reductions in mtDNA:nucDNA copy number persist throughout adulthood. However, mtDNA replication appears to cease between 3 and 6 days of age in untreated nematodes (Rooney et al., 2014). Therefore, it is possible that replication of some mtDNAs may be blocked by lesions until the point at which replication stops, resulting in fewer mtDNA copies per cell. Alternately, it is possible that a reduction in mtDNA copy number results from increased removal. Supporting this theory, oxidative damage in cultured mammalian cells induced strand breaks in mtDNA, leading to degradation of mitochondrial genomes (Shokolenko, Venediktova, Bochkareva, Wilson, & Alexeyev, 2009).

A limitation of our study is the inability to draw conclusions about cell-type specific effects of either mtDNA damage or copy number which may be important because mitochondria vary significantly between cell types (Vafai & Mootha, 2012). Cell- and tissue-specific differences include turnover rates (Menzies & Gold, 1971) and levels of mitophagy (Van Laar et al., 2011), such that damaged genomes or alterations in copy number may persist only in some cell types. This is an intriguing hypothesis, as mitochondrial dysfunction specifically in neurons results in whole organism effects in *C. elegans* (Durieux et al., 2011), and is implicated in many human diseases (Coskun et al., 2012).

## Conclusions

Overall, these results identify life-long redox signaling-mediated alterations in mitochondrial function resulting from a low-level exposure that had no effect on lifespan. The outcomes later in life were largely deleterious, although we cannot exclude that the metabolic changes we observed may be advantageous under certain circumstances. These results are potentially important in the context of human health, because environmental exposures that results in irreparable mtDNA damage are common (Meyer et al., 2013), there are increasing reports of pollutants affecting mitochondria (Meyer & Chan, 2017; Meyer et al., 2018), and we now understand that pollution is the major environmental driver of loss of life globally (Landrigan et al., 2018), despite the fact that we know very little about the toxic effects of the great majority of anthropogenically-produced chemicals (Gross & Birnbaum, 2017).

## Methods

### *C. elegans* Strains and Culture Conditions

Populations of *C. elegans* were maintained on K-agar plates seeded with *E. coli* OP50 bacteria, unless otherwise noted. N2 (wild-type), JK1107 *glp-1(q224)*, *sod-2*(gk257 I), *sod-3* (gk235), *gas-1*(fc21), *mev-1*(Kn1), *glp-1*(e2141), *daf-16*(mu86), *skn-1*(zu670), SJ4100 zclS13[*hsp-6*::GFP], SJ4058 zclS9[*hsp-60*::GFP], and *pkc-1*(nj1) were obtained from the *Caenorhabditis* Genetics Center (CGC), University of Minnesota. PE255 *glp-4(bn2)* were provided by Christina Lagido, University of Aberdeen (Aberdeen, UK). The *sod-2;sod-3* [*sod-2*(gk257) I; *sod-3* (tm760) X] double mutant strain and *sod-1;sod-4;sod-5* triple mutant strain [*sod-1*(tm776) ;*sod-5*(tm1146) II; *sod-4*(gk101) III] were a kind gift from Bart Braeckman (Ghent University, Ghent, Belgium).

### UVC Exposure, DNA damage and genome copy number

UVC exposures were conducted in a custom-built exposure cabinet as previously described (Bess et al., 2012). UVC output was measured with a UVX radiometer (UVP, Upland, CA). Synchronized (by bleach/NaOH egg isolation: (Lewis & Fleming, 1995)). L1 nematodes were maintained on plates that contain no bacterial food, and therefore do not develop for UVC exposures, and were then transferred to seeded plates, as previously described (Leung et al., 2013). Mitochondrial and nuclear DNA damage levels and mtDNA copy number were measured as previously described (Gonzalez-Hunt et al., 2016). All PCR conditions and primer sequences can be found in **Table S1**.

### ATP Levels and Oxygen Consumption

ATP levels were measured in two different strains by two methods. First, using the JK1107 *glp-1(q224)* strain, ATP levels were measured as described in (Brys, Castelein, Matthijssens, Vanfleteren, & Braeckman, 2010) using the Molecular Probes ATP determination Kit (Invitrogen/Life Technologies, Carlsbad, CA, USA). Second, the firefly luciferase expressing PE255 *glp-4(bn2)* strain was used to investigate relative, steady state ATP levels *in vivo*, in live nematodes at 2, 4, 6, 8, 10 and 12 days post final UVC dose, as previously described (Lagido, Pettitt, Flett, & Glover, 2008). Details of both protocols are available in the supplemental methods section. Oxygen consumption was measured using a Seahorse Biosciences XF^e^24 extracellular flux analyzer as described (A. L. Luz et al., 2015).

### Gene Expression Assays

RNA was isolated from between 1000 and 2000 nematodes according to the Qiagen RNeasy Min Kit protocol. cDNA was created from 100 ng of the isolated RNA using the High Capacity Reverse Transcription kit (Life Technologies) per the manufacturer’s instructions.

Gene expression was measured via real-time PCR using the Power SYBR Green PCR Master Mix (Life Technologies). Changes in expression levels were calculated based on the standard delta-delta-cT method, compared to housekeeping genes *cdc-42* and *pmp-3*. Primer sequences and PCR conditions can be found in **Table S2**.

### Targeted Metabolomics

Nematodes (approximately 7500 for each sample) were washed, incubated in K-media at 20°C for 30 minutes to allow for gut clearance, and washed twice in ice cold PBS. Worms were resuspended in 0.6% formic acid and stored at −80°C. Samples were thawed on ice, lysed by sonication and aliquots were removed for total protein determination. 270 uL acetonitrile was added to each sample, and they were vortexed for 1 minute and centrifuged at 15,000 x g for 10 minutes to pellet proteins. Amino acids, acylcarnitines and organic acids were analyzed using stable isotope dilution technique. Amino acids and acylcarnitine measurements were made by flow injection tandem mass spectrometry using sample preparation methods described previously (J. An et al., 2004; Ferrara et al., 2008). The data were acquired using a Waters TQD mass spectrometer equipped with AcquityTM UPLC system and controlled by MassLynx 4.1 operating system (Waters, Milford, MA). Organic acids were quantified using methods described previously (Jensen et al., 2006) employing Trace Ultra GC coupled to ISQ MS operating under Xcalibur 2.2 (Thermo Fisher Scientific, Austin, TX). Metabolite levels were normalized to total protein as determined by BCA (Thermo Scientific, Rockford, IL) and fold changes compared to day 4 control samples were calculated.

### Mitochondrial Isolation and Oxyblot

Mitochondria were isolated essentially as described (Falk, Kayser, Morgan, & Sedensky, 2006). Briefly, worms were washed off plates, rinsed twice in K-media and allowed to clear guts for 30 mins in in K-media at room temperature. All steps from here were conducted on ice.

Worms were washed once in K-media and twice in 10 mL MSM-E buffer (MSM-E – 150 ml MSM (=40.08g mannitol, 23.96g sucrose, 1.047g MOPS, 1L milliQ H2O, pH 7.4,) plus 3mL 0.1M EDTA), resuspended in 1 mL MSM-E and lysed in a glass-Teflon potter homogenizer. Worms were microscopically monitored for lysis. Once sufficiently lysed, one volume MSM-EB (50ml MSM-E + 0.2g Fatty acid free BSA) was added and samples were centrifuged at 300 x G for 10 minutes. Supernatant was transferred to a fresh tube and kept, and pellet was re-extracted. Supernatants from the two extractions were combined and spun at 7000 x G for 10 minutes to yield the mitochondrial pellet, and supernatant samples were taken as the cytoplasmic fraction. Mitochondrial pellets were washed once in MSM-E and once in MSM. Oxyblot was performed following manufacturer’s instructions (MilliporeSigma, Burlington, MA).

### Growth, antioxidant rescue, protein translation exacerbation, and lethality assays

Laval growth was assessed by measuring nematode size at 72 h post-exposure, using either a COPAS Biosorter (Union Biometrica, Holliston, MA), which measures extinction and time of flight of individual *C. elegans* (X. Yang et al., 2012), or microscopic imaging. For the latter, aliquots of nematodes were frozen, thawed and imaged at 10x magnification on a Zeis Axioskop. Nematode length was measured using NIS elements BR software (Nikon Inc. Melville, NY, USA). In many cases, growth is presented as percent control to permit comparisons between the methods and between strains that may develop at different rates under control conditions. In some experiments, larvae were staged (L1 through adult) at 24, 48, 72, and 96 h post-final exposure. For chemical exacerbation experiments, mitochondrial translation inhibitors doxycycline and chloramphenicol were added after plates cooled, and nematodes were exposed to chemicals throughout development after UVC exposure. For trolox, MitoQ, and N-acetyl cysteine antioxidant rescue experiments, compounds were added to the medium prior to pouring plates, nematodes were exposed to antioxidants throughout development after UVC exposure, and size was measured on the COPAS Biosorter. Lethality assays were conducted 4 days after the 3^rd^ UVC exposure. 10-15 nematodes were transferred to plates containing either rotenone or paraquat that were seeded with UVC-inactivated UvrA deficient OP50 *E. coli* to reduce bacterial metabolism of the chemicals (Maurer, Ryde, Yang, & Meyer, 2015). Nematodes were scored for survival 24 hours later and were counted as dead if they did not respond (move) to gentle prodding with a worm pick.

### Lifespan analysis

Lifespan was determined on K agar plates with an OP50 lawn at 20º C. Approximately 25 synchronized (as above) L1 nematodes were plated per condition were assayed in triplicate experiments. Beginning one day after reaching L4 and continuing until reproduction stopped, all adults were transferred daily onto a new plate, leaving offspring on the old plate. Nematodes were monitored daily by tapping the adults on the head. Animals were considered dead if no movement was observed following repeated probing. Individual lifespans were calculated from egg (day 0) until death.

### Analysis of movement

Synchronized *glp-1* L1 larvae were obtained by bleach-sodium hydroxide isolation of eggs (described above), transferred to unseeded no-peptone K-agar, and exposed to 3 sequential doses, 24 hours apart, of 0, 3, or 4 J/m^2^ UVC as described above. Next, nematodes were transferred to K-agar plates seeded with OP50 and were grown at 25° for either 4 or 8 days. For each treatment group, six 60-second videos of nematode locomotion were recorded on a Nikon SMZ1500 stereomicroscope using NIS-Elements BR software. Analysis was completed using Fiji/ImageJ. Nematode pathways were constructed using the first 10 seconds of each video and quantified using the Analyze Particles function. Pathway area was normalized to mean nematode size for each treatment group.

### Measurement of mitochondrial unfolded protein response

UPR^mt^ activation was measured in Hsp-6 (strain SJ4100) and Hsp-60 (strain SJ4058) GFP fusion reporter strain 4 days post 2.5 or 5 J/m^2^ UVC exposure. Chloramphenicol exposure was included as a positive control. Worms were washed with K-medium, and approximately 100 worms in 100 µL K-medium were aliquoted into wells of a white 96 well plate (70 worms per well for SW4100 chloramphenicol exposure). GFP fluorescence was measured from 4 wells on a FLUOstar Optima microplate reader and adjusted for number of worms per well.

### Statistical analysis

Specific analyses are indicated in the manuscript text of figure legends. In Figures, error bars indicate standard errors of the mean. In any case where asterisks are employed, p ≤ 0.05. Correction for multiple comparisons is indicated in the figure legends.

## Acknowledgements

This research was supported by the National Institute of Environmental Health Sciences (R01ES017540, T32ES021432, and P42ES010356). The content is solely the responsibility of the authors and does not necessarily represent the official views of the NIH. We thank Tracey Crocker for assistance with the doxycycline and chloramphenicol experiments.

## Author contributions

John P. Rooney: conceptualization, methodology, formal analysis, investigation, writing - original draft preparation. Kathleen A. Hershberger: formal analysis, writing - original draft preparation, writing - review and editing, visualization. Elena A. Turner: investigation. Lauren J. Donoghue: investigation. Laura L. Maurer: investigation. Ian T. Ryde: investigation. Jina J Kim: investigation. Rashmi Joglekar: investigation. Jonathan D. Hibshman: investigation, writing - review and editing. Latasha L. Smith: investigation. Dhaval P. Bhatt: investigation. Olga R. Ilkayeva: methodology, investigation, resources. Matthew D. Hirschey: supervision. Joel N. Meyer: conceptualization, writing - review and editing, supervision, project administration, funding acquisition.

## References

Agarwal, S., & Ganesh, S. (2020). Perinuclear mitochondrial clustering, increased ROS levels, and HIF1 are required for the activation of HSF1 by heat stress. Journal of Cell Science, 133(13), jcs245589. doi:10.1242/jcs.245589

An, J., Muoio, D. M., Shiota, M., Fujimoto, Y., Cline, G. W., Shulman, G. I., … Newgard, C. B. (2004). Hepatic expression of malonyl-CoA decarboxylase reverses muscle, liver and whole-animal insulin resistance. Nat Med, 10(3), 268–274. doi:10.1038/nm995

An, J. H., & Blackwell, T. K. (2003). SKN-1 links C. elegans mesendodermal specification to a conserved oxidative stress response. Genes Dev, 17(15), 1882–1893. Retrieved from http://www.ncbi.nlm.nih.gov/entrez/query.fcgi?cmd=Retrieve&db=PubMed&dopt=Citation&list_uids=12869585

Apfeld, J., O’Connor, G., McDonagh, T., DiStefano, P. S., & Curtis, R. (2004). The AMP-activated protein kinase AAK-2 links energy levels and insulin-like signals to lifespan in C. elegans. Genes Dev, 18(24), 3004–3009. doi:10.1101/gad.1255404

Barouki, R., Gluckman, P. D., Grandjean, P., Hanson, M., & Heindel, J. J. (2012). Developmental origins of non-communicable disease: implications for research and public health. Environ Health, 11, 42. doi:10.1186/1476-069X-11-42

Benedetti, M. G., Foster, A. L., Vantipalli, M. C., White, M. P., Sampayo, J. N., Gill, M. S., … Lithgow, G. J. (2008). Compounds that confer thermal stress resistance and extended lifespan. Experimental Gerontology, 43(10), 882–891. doi:10.1016/j.exger.2008.08.049

Berthiaume, J. M., & Wallace, K. B. (2007). Adriamycin-induced oxidative mitochondrial cardiotoxicity. Cell Biol Toxicol, 23(1), 15–25. doi:10.1007/s10565-006-0140-y

Bess, A. S., Crocker, T. L., Ryde, I. T., & Meyer, J. N. (2012). Mitochondrial dynamics and autophagy aid in removal of persistent mitochondrial DNA damage in Caenorhabditis elegans. Nucleic Acids Res. doi:10.1093/nar/gks532

Bess, A. S., Ryde, I. T., Hinton, D. E., & Meyer, J. N. (2013). UVC-induced mitochondrial degradation via autophagy correlates with mtDNA damage removal in primary human fibroblasts. J Biochem Mol Toxicol, 27(1), 28–41. doi:10.1002/jbt.21440

Blajszczak, C., & Bonini, M. G. (2017). Mitochondria targeting by environmental stressors: Implications for redox cellular signaling. Toxicology, 391, 84–89. doi:10.1016/j.tox.2017.07.013

Braeckman, B. P., Houthoofd, K., De Vreese, A., & Vanfleteren, J. R. (2002). Assaying metabolic activity in ageing *Caenorhabditis elegans*. Mechanisms of Ageing and Development, 123(2-3), 105–119. Retrieved from <Go to ISI>://000177152000008

Braun, J. M., & Gray, K. (2017). Challenges to studying the health effects of early life environmental chemical exposures on children’s health. PLoS Biol, 15(12), e2002800. doi:10.1371/journal.pbio.2002800

Brys, K., Castelein, N., Matthijssens, F., Vanfleteren, J. R., & Braeckman, B. P. (2010). Disruption of insulin signalling preserves bioenergetic competence of mitochondria in ageing Caenorhabditis elegans. BMC Biol, 8, 91. doi:1741-7007-8-91 [pii] 10.1186/1741-7007-8-91

Calabrese, E. J. (2013). Hormetic mechanisms. Crit Rev Toxicol, 43(7), 580–606. doi:10.3109/10408444.2013.808172

Chinnery, P. F. (2015). Mitochondrial disease in adults: what’s old and what’s new? EMBO Mol Med, 7(12), 1503–1512. doi:10.15252/emmm.201505079

Cline, S. D. (2012). Mitochondrial DNA damage and its consequences for mitochondrial gene expression. Biochim Biophys Acta, 1819(9-10), 979–991. doi:10.1016/j.bbagrm.2012.06.002

Coskun, P., Wyrembak, J., Schriner, S. E., Chen, H. W., Marciniack, C., Laferla, F., & Wallace, D. C. (2012). A mitochondrial etiology of Alzheimer and Parkinson disease. Biochim Biophys Acta, 1820(5), 553–564. doi:10.1016/j.bbagen.2011.08.008

Dillin, A., Hsu, A. L., Arantes-Oliveira, N., Lehrer-Graiwer, J., Hsin, H., Fraser, A. G., … Kenyon, C. (2002). Rates of behavior and aging specified by mitochondrial function during development. Science, 298(5602), 2398–2401. Retrieved from http://www.ncbi.nlm.nih.gov/entrez/query.fcgi?cmd=Retrieve&db=PubMed&dopt=Citation&list_uids=12471266

Dingley, S., Polyak, E., Lightfoot, R., Ostrovsky, J., Rao, M., Greco, T., … Falk, M. J. (2010). Mitochondrial respiratory chain dysfunction variably increases oxidant stress in Caenorhabditis elegans. Mitochondrion, 10(2), 125–136. doi:S1567-7249(09)00172-X [pii] 10.1016/j.mito.2009.11.003

Durieux, J., Wolff, S., & Dillin, A. (2011). The cell-non-autonomous nature of electron transport chain-mediated longevity. Cell, 144(1), 79–91. doi:10.1016/j.cell.2010.12.016

Dykens, J. A., & Will, Y. (2007). The significance of mitochondrial toxicity testing in drug development. Drug Discovery Today, 12(17-18), 777–785. doi:DOI 10.1016/j.drudis.2007.07.013

Falk, M. J., Kayser, E. B., Morgan, P. G., & Sedensky, M. M. (2006). Mitochondrial complex I function modulates volatile anesthetic sensitivity in C. elegans. Curr Biol, 16(16), 1641–1645. doi:10.1016/j.cub.2006.06.072

Ferrara, C. T., Wang, P., Neto, E. C., Stevens, R. D., Bain, J. R., Wenner, B. R., … Attie, A. D. (2008). Genetic networks of liver metabolism revealed by integration of metabolic and transcriptional profiling. PLoS Genet, 4(3), e1000034. doi:10.1371/journal.pgen.1000034

Gonzalez-Hunt, C. P., Rooney, J. P., Ryde, I. T., Anbalagan, C., Joglekar, R., & Meyer, J. N. (2016). PCR-Based Analysis of Mitochondrial DNA Copy Number, Mitochondrial DNA Damage, and Nuclear DNA Damage. Curr Protoc Toxicol, 67, 20 11 21–20 11 25. doi:10.1002/0471140856.tx2011s67

Grandjean, P., Barouki, R., Bellinger, D. C., Casteleyn, L., Chadwick, L. H., Cordier, S., … Heindel, J. J. (2015). Life-Long Implications of Developmental Exposure to Environmental Stressors: New Perspectives. Endocrinology, 156(10), 3408–3415. doi:10.1210/EN.2015-1350

Graziewicz, M. A., Sayer, J. M., Jerina, D. M., & Copeland, W. C. (2004). Nucleotide incorporation by human DNA polymerase gamma opposite benzo[a]pyrene and benzo[c]phenanthrene diol epoxide adducts of deoxyguanosine and deoxyadenosine. Nucleic Acids Res, 32(1), 397–405. Retrieved from http://www.ncbi.nlm.nih.gov/entrez/query.fcgi?cmd=Retrieve&db=PubMed&dopt=Citation&list_uids=14729924

Gross, L., & Birnbaum, L. S. (2017). Regulating toxic chemicals for public and environmental health. PLoS Biol, 15(12), e2004814. doi:10.1371/journal.pbio.2004814

Gruber, J., Ng, L. F., Fong, S., Wong, Y. T., Koh, S. A., Chen, C. B., … Halliwell, B. (2011). Mitochondrial changes in ageing Caenorhabditis elegans--what do we learn from superoxide dismutase knockouts? PLoS ONE, 6(5), e19444. doi:10.1371/journal.pone.0019444

Hayes, J. D., & Dinkova-Kostova, A. T. (2014). The Nrf2 regulatory network provides an interface between redox and intermediary metabolism. Trends Biochem Sci, 39(4), 199–218. doi:10.1016/j.tibs.2014.02.002

Hernandez-Garcia, D., Wood, C. D., Castro-Obregon, S., & Covarrubias, L. (2010). Reactive oxygen species: A radical role in development? Free Radic Biol Med, 49(2), 130–143. doi:10.1016/j.freeradbiomed.2010.03.020

Houtkooper, R. H., Mouchiroud, L., Ryu, D., Moullan, N., Katsyuba, E., Knott, G., … Auwerx, J. (2013). Mitonuclear protein imbalance as a conserved longevity mechanism. Nature, 497(7450), 451–457. doi:10.1038/nature12188

Hudson, M. M., Ness, K. K., Gurney, J. G., Mulrooney, D. A., Chemaitilly, W., Krull, K. R., … Robison, L. L. (2013). Clinical ascertainment of health outcomes among adults treated for childhood cancer. JAMA, 309(22), 2371–2381. doi:10.1001/jama.2013.6296

Jensen, M. V., Joseph, J. W., Ilkayeva, O., Burgess, S., Lu, D., Ronnebaum, S. M., … Newgard, C. B. (2006). Compensatory responses to pyruvate carboxylase suppression in islet beta-cells. Preservation of glucose-stimulated insulin secretion. J Biol Chem, 281(31), 22342–22351. doi:10.1074/jbc.M604350200

Kasiviswanathan, R., Gustafson, M. A., Copeland, W. C., & Meyer, J. N. (2012). Human mitochondrial DNA polymerase gamma exhibits potential for bypass and mutagenesis at UV-induced cyclobutane thymine dimers. J Biol Chem, 287(12), 9222–9229. doi:10.1074/jbc.M111.306852

Kimble, J., & Crittenden, S. L. (2005). Germline proliferation and its control. WormBook, 1–14. Retrieved from http://www.ncbi.nlm.nih.gov/entrez/query.fcgi?cmd=Retrieve&db=PubMed&dopt=Citation&list_uids=18050413

Kumsta, C., Thamsen, M., & Jakob, U. (2011). Effects of oxidative stress on behavior, physiology, and the redox thiol proteome of Caenorhabditis elegans. Antioxid Redox Signal, 14(6), 1023–1037. doi:10.1089/ars.2010.3203

Labbadia, J., Brielmann, R. M., Neto, M. F., Lin, Y. F., Haynes, C. M., & Morimoto, R. I. (2017). Mitochondrial Stress Restores the Heat Shock Response and Prevents Proteostasis Collapse during Aging. Cell Rep, 21(6), 1481–1494. doi:10.1016/j.celrep.2017.10.038

Lagido, C., Pettitt, J., Flett, A., & Glover, L. A. (2008). Bridging the phenotypic gap: real-time assessment of mitochondrial function and metabolism of the nematode Caenorhabditis elegans. BMC Physiol, 8, 7. Retrieved from http://www.ncbi.nlm.nih.gov/entrez/query.fcgi?cmd=Retrieve&db=PubMed&dopt=Citation&list_uids=18384668

Landrigan, P. J., Fuller, R., Acosta, N. J. R., Adeyi, O., Arnold, R., Basu, N. N., … Zhong, M. (2018). The Lancet Commission on pollution and health. Lancet, 391(10119), 462–512. doi:10.1016/S0140-6736(17)32345-0

Leung, M. C., Rooney, J. P., Ryde, I. T., Bernal, A. J., Bess, A. S., Crocker, T. L., … Meyer, J. N. (2013). Effects of early life exposure to ultraviolet C radiation on mitochondrial DNA content, transcription, ATP production, and oxygen consumption in developing Caenorhabditis elegans. BMC Pharmacol Toxicol, 14, 9. doi:10.1186/2050-6511-14-9

Lewis, J. A., & Fleming, J. T. (1995). Basic Culture Methods. In H. F. Epstein & D. C. Shakes (Eds.), Caenorhabditis elegans: Modern Biological Analysis of an Organism (pp. 3–29). San Digo, CA: Academic Press.

Lin, K., Dorman, J. B., Rodan, A., & Kenyon, C. (1997). daf-16: An HNF-3/forkhead family member that can function to double the life-span of Caenorhabditis elegans. Science, 278(5341), 1319–1322. Retrieved from https://www.ncbi.nlm.nih.gov/pubmed/9360933

Liu, Y., Nguyen, P., Baris, T. Z., & Poirier, M. C. (2012). Molecular analysis of mitochondrial compromise in rodent cardiomyocytes exposed long term to nucleoside reverse transcriptase inhibitors (NRTIs). Cardiovasc Toxicol, 12(2), 123–134. doi:10.1007/s12012-011-9148-5

Liu, Y., Shim Park, E., Gibbons, A. T., Shide, E. D., Divi, R. L., Woodward, R. A., & Poirier, M. C. (2016). Mitochondrial compromise in 3-year old patas monkeys exposed in utero to human-equivalent antiretroviral therapies. Environ Mol Mutagen, 57(7), 526–534. doi:10.1002/em.22033

Lozoya, O. A., Xu, F., Grenet, D., Wang, T., Grimm, S. A., Godfrey, V., … Santos, J. H. (2020). Single Nucleotide Resolution Analysis Reveals Pervasive, Long-Lasting DNA Methylation Changes by Developmental Exposure to a Mitochondrial Toxicant. Cell Rep, 32(11), 108131. doi:10.1016/j.celrep.2020.108131

Luz, A. L., Godebo, T. R., Smith, L. L., Leuthner, T. C., Maurer, L. L., & Meyer, J. N. (2017). Deficiencies in mitochondrial dynamics sensitize Caenorhabditis elegans to arsenite and other mitochondrial toxicants by reducing mitochondrial adaptability. Toxicology, 387, 81–94. doi:10.1016/j.tox.2017.05.018

Luz, A. L., & Meyer, J. N. (2016). Effects of reduced mitochondrial DNA content on secondary mitochondrial toxicant exposure in Caenorhabditis elegans. Mitochondrion, 30, 255–264. doi:10.1016/j.mito.2016.08.014

Luz, A. L., Rooney, J. P., Kubik, L. L., Gonzalez, C. P., Song, D. H., & Meyer, J. N. (2015). Mitochondrial Morphology and Fundamental Parameters of the Mitochondrial Respiratory Chain Are Altered in Caenorhabditis elegans Strains Deficient in Mitochondrial Dynamics and Homeostasis Processes. PLoS ONE, 10(6), e0130940. doi:10.1371/journal.pone.0130940

Luz, A. L., Smith, L. L., Rooney, J. P., & Meyer, J. N. (2015). Seahorse Extracellular Flux-based analysis of cellular respiration in Caenorhabditis elegans. Current Protocols in Toxicology, 66, 25.27.21–25.27.15.

Maurer, L. L., Ryde, I. T., Yang, X., & Meyer, J. N. (2015). Caenorhabditis elegans as a Model for Toxic Effects of Nanoparticles: Lethality, Growth, and Reproduction. Curr Protoc Toxicol, 66, 20 10 21–20 10 25. doi:10.1002/0471140856.tx2010s66

McCord, J. M., & Fridovich, I. (1969). Superoxide dismutase. An enzymic function for erythrocuprein (hemocuprein). J Biol Chem, 244(22), 6049–6055. Retrieved from https://www.ncbi.nlm.nih.gov/pubmed/5389100

Menzies, R. A., & Gold, P. H. (1971). The turnover of mitochondria in a variety of tissues of young adult and aged rats. J Biol Chem, 246(8), 2425–2429. Retrieved from https://www.ncbi.nlm.nih.gov/pubmed/5553400

Meredith, M. J., & Reed, D. J. (1982). Status of the mitochondrial pool of glutathione in the isolated hepatocyte. J Biol Chem, 257(7), 3747–3753. Retrieved from https://www.ncbi.nlm.nih.gov/pubmed/7061508

Meyer, J. N., & Chan, S. S. L. (2017). Sources, mechanisms, and consequences of chemical-induced mitochondrial toxicity. Toxicology. doi:10.1016/j.tox.2017.06.002

Meyer, J. N., Hartman, J. H., & Mello, D. F. (2018). Mitochondrial Toxicity. Toxicol Sci, 162(1), 15–23. doi:10.1093/toxsci/kfy008

Meyer, J. N., Leung, M. C., Rooney, J. P., Sendoel, A., Hengartner, M. O., Kisby, G. E., & Bess, A. S. (2013). Mitochondria as a target of environmental toxicants. Toxicol Sci, 134(1), 1–17. doi:10.1093/toxsci/kft102

Mouchiroud, L., Houtkooper, R. H., Moullan, N., Katsyuba, E., Ryu, D., Canto, C., … Auwerx, J. (2013). The NAD(+)/Sirtuin Pathway Modulates Longevity through Activation of Mitochondrial UPR and FOXO Signaling. Cell, 154(2), 430–441. doi:10.1016/j.cell.2013.06.016

Murphy, C. T., McCarroll, S. A., Bargmann, C. I., Fraser, A., Kamath, R. S., Ahringer, J., … Kenyon, C. (2003). Genes that act downstream of DAF-16 to influence the lifespan of Caenorhabditis elegans. Nature, 424(6946), 277–283. Retrieved from http://www.ncbi.nlm.nih.gov/entrez/query.fcgi?cmd=Retrieve&db=PubMed&dopt=Citation&list_uids=12845331

Murphy, M. P. (2009). How mitochondria produce reactive oxygen species. Biochem J, 417(1), 1–13. doi:10.1042/BJ20081386

Ng, L. F., Ng, L. T., van Breugel, M., Halliwell, B., & Gruber, J. (2019). Mitochondrial DNA Damage Does Not Determine C. elegans Lifespan. Front Genet, 10, 311. doi:10.3389/fgene.2019.00311

Nunnari, J., & Suomalainen, A. (2012). Mitochondria: in sickness and in health. Cell, 148(6), 1145–1159. doi:10.1016/j.cell.2012.02.035

Poirier, M. C., Gibbons, A. T., Rugeles, M. T., Andre-Schmutz, I., & Blanche, S. (2015). Fetal consequences of maternal antiretroviral nucleoside reverse transcriptase inhibitor use in human and nonhuman primate pregnancy. Curr Opin Pediatr, 27(2), 233–239. doi:10.1097/MOP.0000000000000193

Rauthan, M., & Pilon, M. (2015). A chemical screen to identify inducers of the mitochondrial unfolded protein response in C. elegans. Worm, 4(4), e1096490. doi:10.1080/21624054.2015.1096490

Rea, S. L., Ventura, N., & Johnson, T. E. (2007). Relationship between mitochondrial electron transport chain dysfunction, development, and life extension in Caenorhabditis elegans. PLoS Biol, 5(10), e259. Retrieved from http://www.ncbi.nlm.nih.gov/entrez/query.fcgi?cmd=Retrieve&db=PubMed&dopt=Citation&list_uids=17914900

Rooney, J. P., Luz, A. L., Gonzalez-Hunt, C. P., Bodhicharla, R., Ryde, I. T., Anbalagan, C., & Meyer, J. N. (2014). Effects of 5 ’-fluoro-2-deoxyuridine on mitochondrial biology in Caenorhabditis elegans. Experimental Gerontology, 56, 69–76. doi:DOI 10.1016/j.exger.2014.03.021

Roubicek, D. A., & de Souza-Pinto, N. C. (2017). Mitochondria and mitochondrial DNA as relevant targets for environmental contaminants. Toxicology, 391, 100–108. doi:10.1016/j.tox.2017.06.012

Shokolenko, I., Venediktova, N., Bochkareva, A., Wilson, G. L., & Alexeyev, M. F. (2009). Oxidative stress induces degradation of mitochondrial DNA. Nucleic Acids Res, 37(8), 2539–2548. doi:Doi 10.1093/Nar/Gkp100

Shpilka, T., & Haynes, C. M. (2018). The mitochondrial UPR: mechanisms, physiological functions and implications in ageing. Nat Rev Mol Cell Biol, 19(2), 109–120. doi:10.1038/nrm.2017.110

Suomalainen, A., & Isohanni, P. (2010). Mitochondrial DNA depletion syndromes--many genes, common mechanisms. Neuromuscular disorders : NMD, 20(7), 429–437. doi:10.1016/j.nmd.2010.03.017

Tsang, W. Y., & Lemire, B. D. (2002). Mitochondrial genome content is regulated during nematode development. Biochem Biophys Res Commun, 291(1), 8–16. Retrieved from http://www.ncbi.nlm.nih.gov/entrez/query.fcgi?cmd=Retrieve&db=PubMed&dopt=Citation&list_uids=11829454

Tsang, W. Y., Sayles, L. C., Grad, L. I., Pilgrim, D. B., & Lemire, B. D. (2001). Mitochondrial respiratory chain deficiency in Caenorhabditis elegans results in developmental arrest and increased life span. J Biol Chem, 276(34), 32240–32246. Retrieved from http://www.ncbi.nlm.nih.gov/entrez/query.fcgi?cmd=Retrieve&db=PubMed&dopt=Citation&list_uids=11410594

Vafai, S. B., & Mootha, V. K. (2012). Mitochondrial disorders as windows into an ancient organelle. Nature, 491(7424), 374–383. doi:10.1038/nature11707

Van Gilst, M. R., Hadjivassiliou, H., Jolly, A., & Yamamoto, K. R. (2005). Nuclear hormone receptor NHR-49 controls fat consumption and fatty acid composition in C. elegans. PLoS Biol, 3(2), e53. doi:10.1371/journal.pbio.0030053

Van Laar, V. S., Arnold, B., Cassady, S. J., Chu, C. T., Burton, E. A., & Berman, S. B. (2011). Bioenergetics of neurons inhibit the translocation response of Parkin following rapid mitochondrial depolarization. Human molecular genetics, 20(5), 927–940. doi:10.1093/hmg/ddq531

Wood, C. E., Hester, S. D., Chorley, B. N., Carswell, G., George, M. H., Ward, W., … Deangelo, A. B. (2015). Latent carcinogenicity of early-life exposure to dichloroacetic acid in mice. Carcinogenesis, 36(7), 782–791. doi:10.1093/carcin/bgv057

Yang, W., & Hekimi, S. (2010). A mitochondrial superoxide signal triggers increased longevity in Caenorhabditis elegans. PLoS Biol, 8(12), e1000556. doi:10.1371/journal.pbio.1000556

Yang, X., Gondikas, A. P., Marinakos, S. M., Auffan, M., Liu, J., Hsu-Kim, H., & Meyer, J. N. (2012). Mechanism of silver nanoparticle toxicity is dependent on dissolved silver and surface coating in Caenorhabditis elegans. Environ Sci Technol, 46(2), 1119–1127. doi:10.1021/es202417t

Yee, C., Yang, W., & Hekimi, S. (2014). The intrinsic apoptosis pathway mediates the pro-longevity response to mitochondrial ROS in C. elegans. Cell, 157(4), 897–909. doi:10.1016/j.cell.2014.02.055

Yun, J., & Finkel, T. (2014). Mitohormesis. Cell Metab, 19(5), 757–766. doi:10.1016/j.cmet.2014.01.011

Zubovych, I. O., Straud, S., & Roth, M. G. (2010). Mitochondrial dysfunction confers resistance to multiple drugs in Caenorhabditis elegans. Mol Biol Cell, 21(6), 956–969. doi:10.1091/mbc.E09-08-0673

