## Supplemental Files for "Early-life mitochondrial DNA damage results in lifelong deficits in energy production mediated by redox signaling in *Caenorhabditis elegans*"

### Supplemental Figures and Tables

John P. Rooney[1]*, Kathleen A. Hershberger[2]*, Elena A. Turner[1], Lauren J. Donoghue[1], Laura L. Maurer[1], Ian T. Ryde[1], Jina J Kim[1], Rashmi Joglekar[1], Jonathan D. Hibshman[3], Latasha L. Smith[1], Dhaval P. Bhatt[4], Olga R. Ilkayeva[4], Matthew D. Hirschey[4], and Joel N. Meyer[1,5]

[1]Duke University, Nicholas School of the Environment, Integrated Toxicology and Environmental Health Program, Durham, NC

[2]Duke University, Nicholas School of the Environment, Durham, NC

[3]Duke University Department of Biology and University Program in Genetics and Genomics, Durham, NC

[4]Duke Molecular Physiology Institute, Durham, NC

[5]Corresponding author.

* These authors contributed equally to this work.

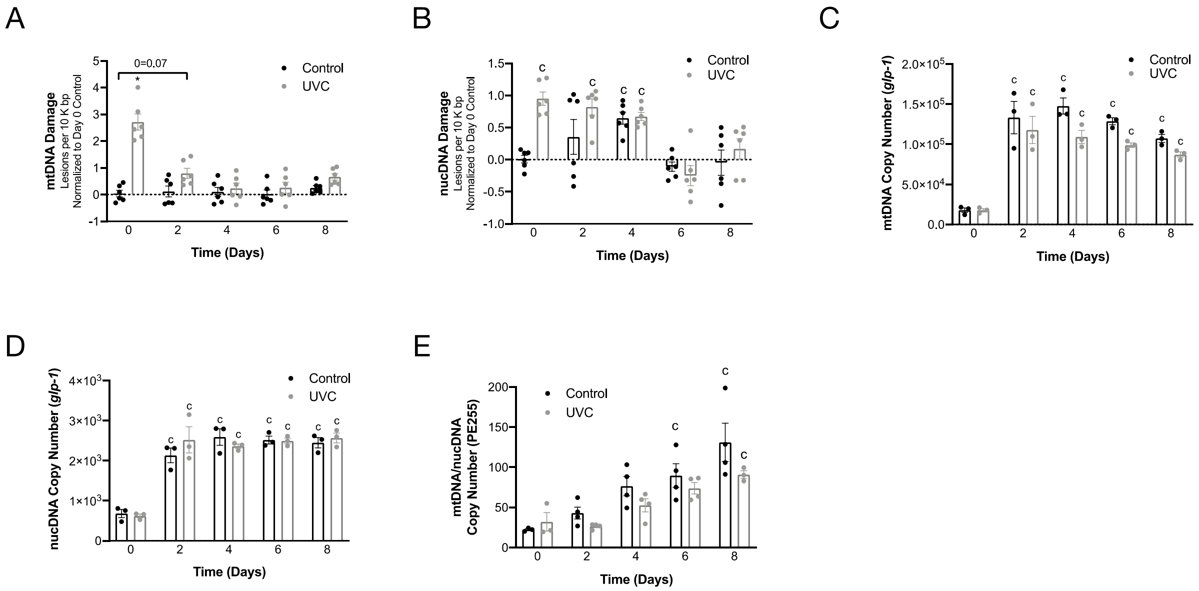


**Figure S1. UVC exposure reduced mtDNA copy number in JK1107 (*glp-1*) and PE255 individuals**. **A.** mtDNA damage data from figure 1C normalized to control day 0. Two-way ANOVA with Bonferroni's correction for multiple comparisons (13 comparisons). n = 3 per group per experiment (2 independent experiments). * $p \leq 0.05$. **B.** nucDNA damage from figure 1D normalized to control day 0. Two-way ANOVA with Bonferroni's correction for multiple comparisons (13 comparisons). n = 3 per group per experiment (2 independent experiments). "c" indicates $p \leq 0.05$ compared to day 0 control**. C.** mtDNA copy number per worm is reduced in response to UVC exposure *in glp*-1 nematodes. Two-way ANOVA with Bonferroni's correction for multiple comparisons (13 comparisons). n = 1 per group per experiment (3 independent experiments). "c" indicates $p \leq 0.05$ compared to day 0 control. Multiple comparisons revealed no significant differences between control and UVC exposed nematodes at individual timepoints, but there was an overall significant effect of UVC exposure (p = 0.0029). **D.** nucDNA copy number is unchanged in *glp-1* nematodes exposed to UVC compared to control nematodes. Two-way ANOVA with Bonferroni's correction for multiple comparisons (13 comparisons). n = 1 per group per experiment (3 independent experiments). "c" indicates $p \leq 0.05$ compared to day 0 control. **E**. mtDNA/nucDNA ratio is reduced in PE255 nematodes. Two-way ANOVA with Bonferroni's correction for multiple comparisons (13 comparisons). n = 1 per group per experiment (3–4 independent experiments). "c" indicates $p \leq 0.05$ compared to day 0 control. Multiple comparisons revealed no significant differences between control and UVC exposed nematodes at individual timepoints, but there was an overall significant effect of UVC exposure (p = 0.031).

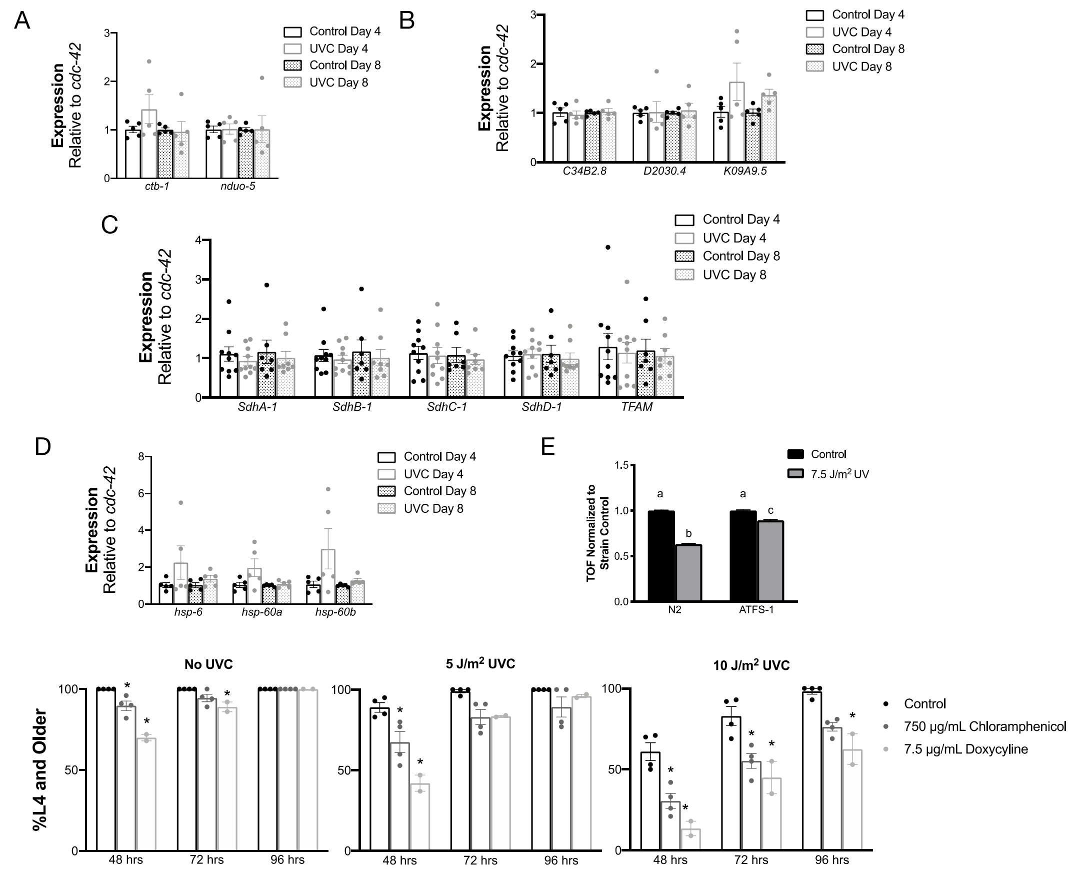


**Figure S2. Transcriptional response to UVC exposure is unremarkable. A.** mtDNA-encoded subunits: *ctb-1* and *nduo-5*. **B.** nucDNA-encoded subunits: C34B2.8, D2030.4, K09A9.5. All are part of complex I of the MRC, except *ctb-1*, which is part of Complex III. n = 5 **C.** Transcript levels of MRC complex II subunits are unchanged in response to UVC exposure. Transcript levels of *hmg-5*, the nematode homologue of Tfam (a structural component of the nucleoid and the mtDNA transcription factor) is also unchanged. n = 7-10. **D.** Transcript levels of mitochondrial chaperone proteins *hsp-6*, *hsp-60a* and *hsp-60b* are unchanged. n = 5. **E.** *atfs-1* mutants are protected, compared to wild-type N2, against UVC-induced developmental delay. UVC exposed TOF values are normalized to control values per strain. n > 2000 for each group. 2-Way ANOVA with Tukey's correction for multiple comparisons. Letters indicate $p \leq 0.05$. **F.** Co-exposure to UVC (5 or 10 J/m$^2$) and mitochondrial translation inhibitors chloramphenicol (750 µg/ml) or doxycycline (7.5 µg/ml) further delays L4 development compared to translation inhibitors alone. Two-way ANOVA with Tukey's correction for multiple comparisons. *$p \leq 0.05$ compared to no treatment control at each time point. One (doxycycline) or two independent experiments, n = 20-30 nematodes per well in each experiment, 2 wells per experiment.

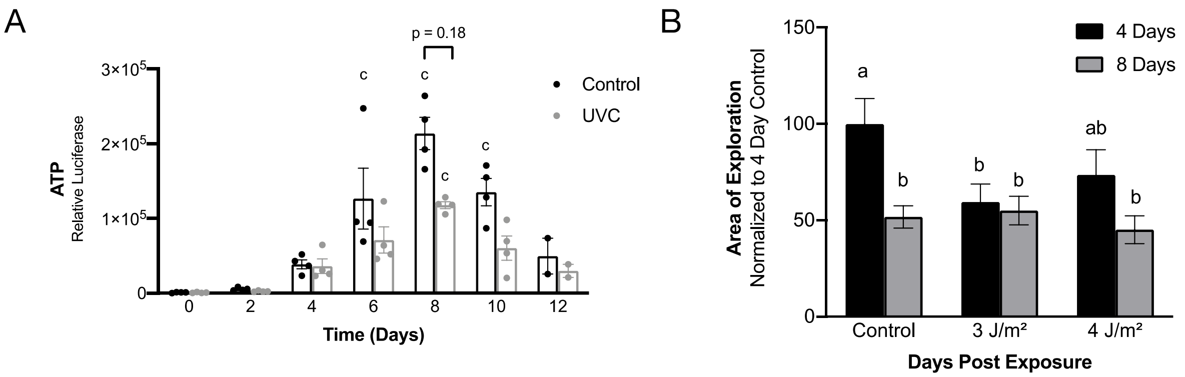


**Figure S3. Decreased energy availability in nematodes exposed to UVC in early life. A.** ATP measurements in the ATP reporter strain PE255 show that ATP levels are reduced throughout life in the UVC exposed animals ($p < 0.0001$), though there were no significant differences between UVC and control animals at specific time points. n = 2-4, from 2-4 independent experiments. 2-Way ANOVA with Bonferroni correction for multiple comparisons (19 comparisons). "c" indicates $p \leq 0.05$ compared to control at day 0. **B.** Area of exploration decreased in worms 8 days post UVC exposure. Letters show which groups are significantly different ($p \leq 0.05$). One-way ANOVA with Tukey's correction for multiple comparisons. n=130–175 individuals in 3 experimental replicates.

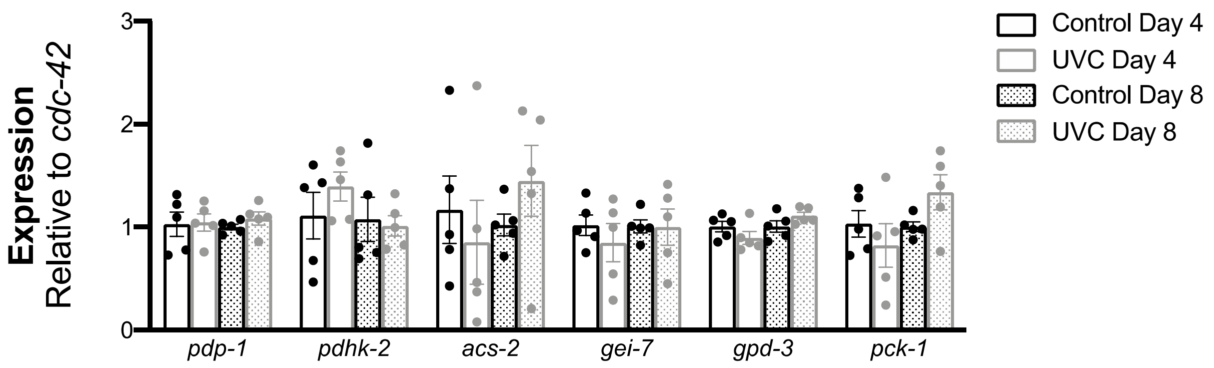


**Figure S4. Transcript levels of metabolic enzymes that regulate TCA cycle metabolism are unchanged in later life with early life UVC exposure.** Transcript levels are unchanged after UVC exposure for: *pdp-1* (pyruvate dehydrogenase phosphatase), *pdhk-2* (pyruvate dehydrogenase kinase), *acs-2* (Acyl-CoA synthetase), *gei-7* (isocitrate lyase/malate synthase), *gpd-3* (glyceraldehyde 3-phosphate dehydrogenase), and *pck-1* (phosphoenolpyruvate carboxykinase). n = 5 per group. Two-Way ANOVA with Tukey correction multiple comparisons.

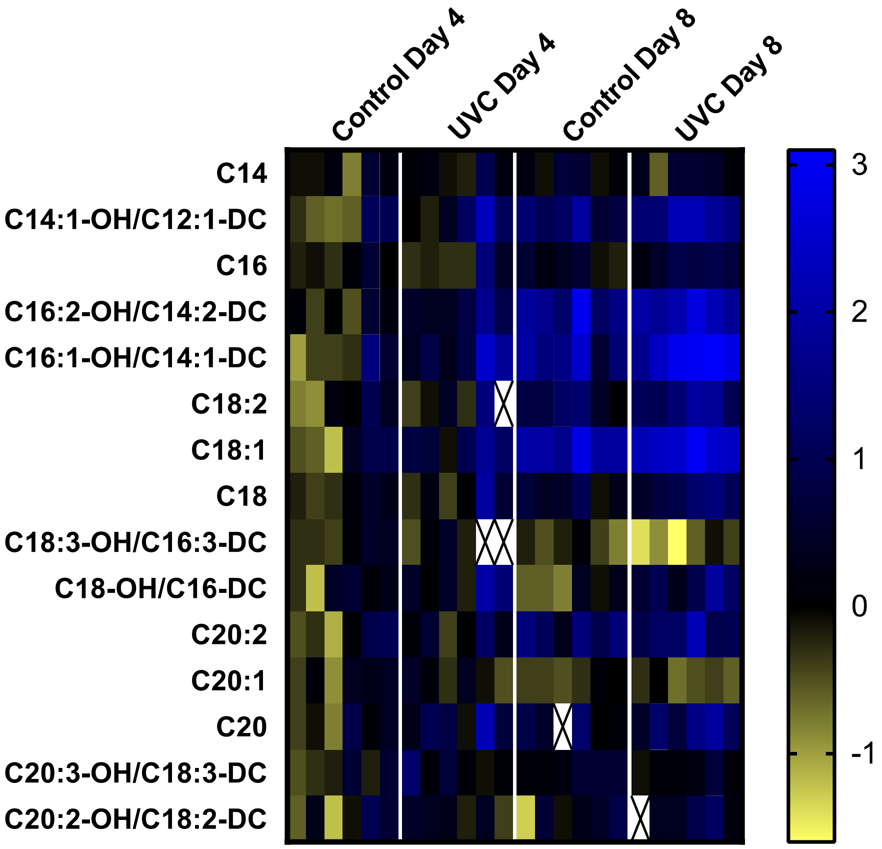


**Figure S5: Elevation of long chain acyl-carnitines in UVC exposed worms.** Heat map of long chain acyl carnitines 4- and 8-days post-exposure in control and UVC exposed animals.

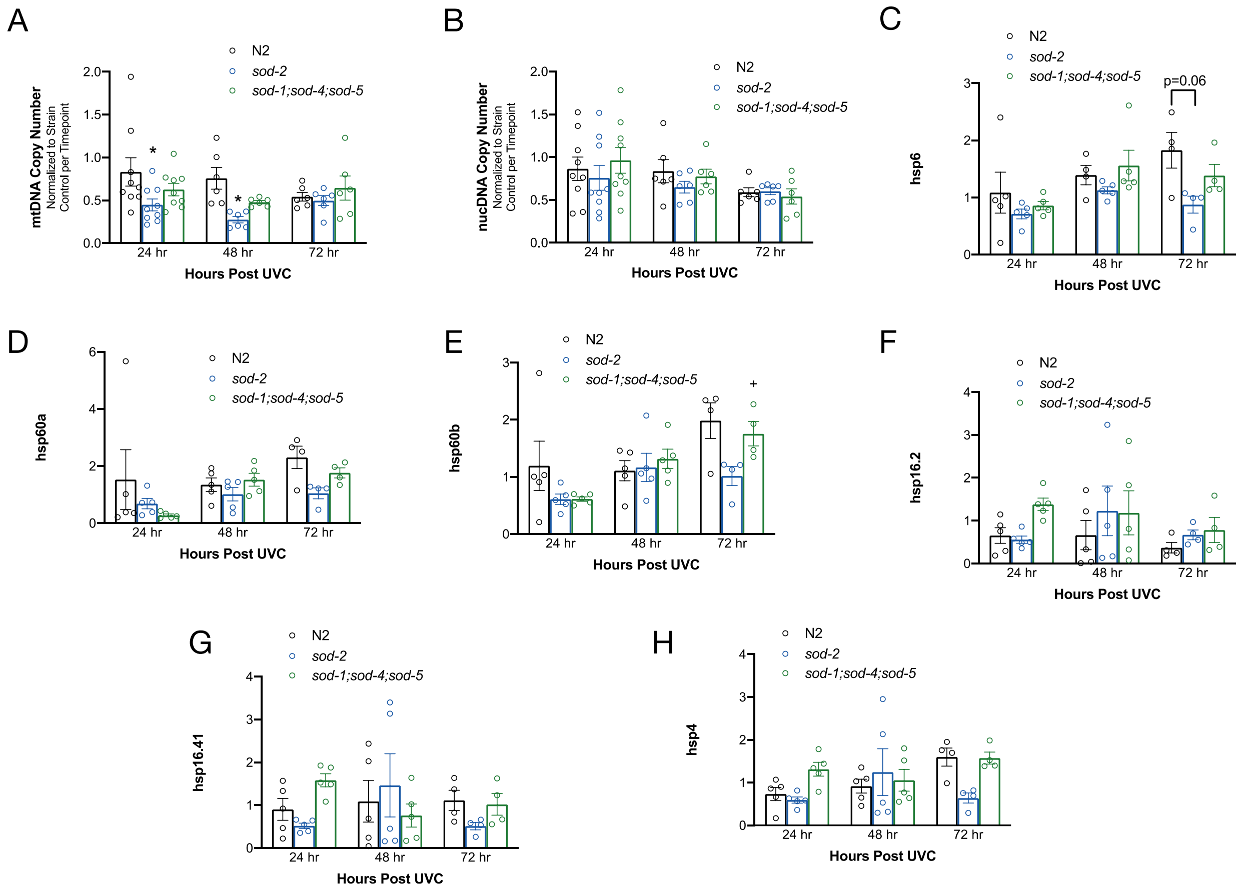


**Figure S6. Characterization of SOD mutants in response to UVC. A**. mtDNA and **B.** nucDNA copy number in SOD mutants over time. Each UVC exposed strain is normalized to its own non-exposed control at each time point. Two-way ANOVA with Bonferroni correction for multiple comparisons (12 comparisons). n = 6–9 per group. * $p \leq 0.05$ compared to N2 UVC exposed at same time point. There is a delay in mitochondrial copy number expansion in the *sod-2* mutant, but no differences in nucDNA copy number. Transcript levels of mitochondrial heat shock proteins: **C.** *hsp-6* **D.** *hsp-60a* and **E.** *hsp-60b*. Transcript levels of cytosolic heat shock proteins: **F.** *hsp-16.12* and G. *hsp-16.41*. **H.** Transcript level of endoplasmic reticular *hsp-4*. Transcript expression is relative to *cdc-42* expression. Two-way ANOVA with Bonferroni correction for multiple comparisons (12 comparisons). n = 4–5 per group.

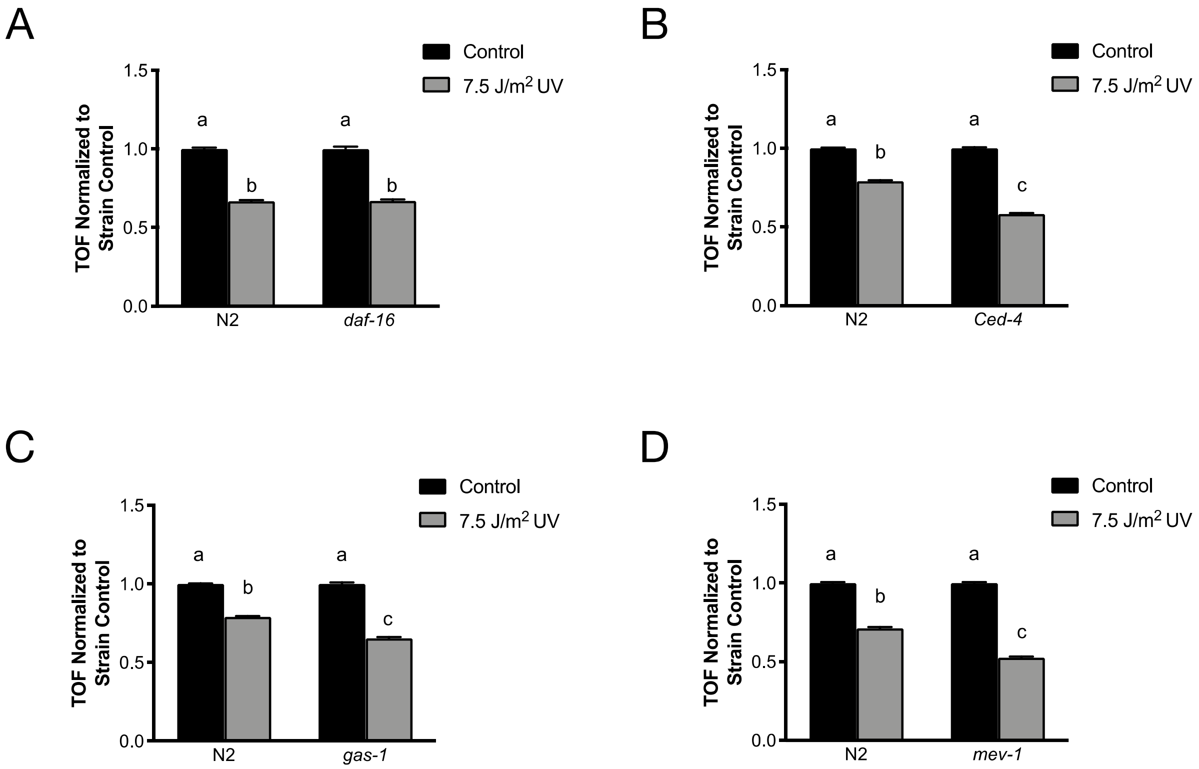


**Figure S7.** ***daf-16, ced-4, gas-1,*** **and** ***mev-1*** **mutants are more sensitive to UVC. A.** *daf-16* mutants show a similar growth delay to UVC compared to N2 controls. **B.** *ced-4* mutants are more sensitive to UVC induced larval growth inhibition than N2. Both **C.** *gas-1* (complex I) and **D.** *mev-1* (complex II) mutants are more significantly growth delayed than wild type in response to UVC exposure. n > 500 in each group from 2 independent experiments. TOF values from UVC exposures are normalized to strain controls. Letters show which groups are significantly different ($p \leq 0.05$). Two-way ANOVA with Tukey’s correction for multiple comparisons.

**Table S1: Primers for DNA Damage and Copy Number Assays in *C. elegans***

| Target | Direction | Sequence (5′-3′) | Annealing Temp. ($^{o}$C) | Ref. |
|---|---|---|---|---|
| mtDNA, long (10.9 kb) | F | CCATCAATTGCCCAAAGGGGAGT | 64 | [1] |
| | R | TGTCCTCAAGGCTACCACCTTCTTCA | | |
| nucDNA, long (9.3 kb) | F | TGGCTGGAACGAACCGAACCAT | 64 | [1] |
| | R | GGC GGT TGT GGA GTG TGG GAA G | | |
| mtDNA, short (75 bps) | F | AGCGTCATTTATTGGGAAGAAGAC | 60 | [2] |
| | R | AAGCTTGTGCTAATCCCATAAATGT | | |
| nucDNA, short (164 bps) | F | GCCGACTGGAAGAACTTGTC | 60 | [3] |
| | R | GCG GAG ATC ACC TTCCAG TA | | |

**Table S2: Real Time PCR Primers for Gene Expression**

| Gene | Direction | 5'-3' Seq | Amplicon Size |
|---|---|---|---|
| acs-2 | F | GGCACACCGACCATGTTTAT | 194 |
| | R | ATGGTGACTAGAGGGGATGTC | |
| C34B2.8 | F | CTTTTCCGAAGCTTGTCTGG | 197 |
| | R | CTTGGCCAACAATTTGAGC | |
| ctb-1 | F | TTCCAATTTGAGGGCCAACT | 116 |
| | R | AACTAGAATAGCTCACGGCAATAAAAA | |
| D2030.4 | F | GCGAGATGAAGGCTACTTGG | 115 |
| | R | GGTGCATTTTGGGTTTGG | |
| K09A9.5 (gas-1) | F | AGTCATCATCAAGGCCATCC | 185 |
| | R | TTGTTGGGATGTCAATACCG | |
| gei-7 (icl-1) | F | TGCTCATCCAGGATTGGTGC | 198 |
| | R | CTGAGCCAAGAGTCGAGGTATCCA | |
| gpd-3 | F | CCAGTACGATTCCACTCACGGA | 104 |
| | R | CTGGGTCTCTTGAGTTGTAGACCTT | |
| hmg-5 (TFAM) | F | TGTCTGGAGCTGGAATGGAA | 108 |
| | R | GCTTCTTCGCTTCGTCTGTG | |
| hsp-16.2 | F | CGCCAAAGAAAGAAGCGGTT | 60 |
| | R | CTTCGACGATTGCCTGTTGA | |
| hsp-16.41 | F | TGGACGAACTCACTGGATCTG | 133 |
| | R | TGAGAGACATCGAGTTGAACCG | |
| hsp-4 | F | CGTTCAAGATCGTCGACAAGT | 138 |
| | R | GACCAAGGTAGGATTCGGCA | |
| hsp-6 | F | TCGTGTCATCAACGAGCCAA | 76 |
| | R | AGCGATGATCTTATCTCCAGCG | |
| hsp-60a | F | AGGCTCTTACCACTCTTGTTCT | 123 |
| | R | CTCCCGTCGCAATTCCCATA | |
| hsp-60b | F | CCAAGAAGGTCACCATCACC | 64 |
| | R | TCTGTTTGATCTCCACGCCC | |
| mev-1 (sdhC-1) | F | GTTGGACAGATCTACAAATCGGG | 100 |
| | R | TCTTGTTGCTCTTGTTCTGGC | |
| nd-5 | F | TTAGCAAGTTTGGTCGAAGAAGATT | 88 |
| | R | GGCCCAAAGTAACTATTGAAAAACC | |
| pck-1 | F | GGGACTTCCACGTCCAGTTAAGCAA | 117 |
| | R | TAGCCCGAGCCGAATGACCA | |
| pdhk-2 | F | CGAAACAATGGCTGAAGGAT | 208 |
| | R | CACATCACAGGCAGGATCAA | |
| pdp-1 | F | CTCACGATGGAATGCTGATG | 109 |

| | | | |
|---|---|---|---|
| | R | TGGATTATTTGCGGCTAGTTG | |
| polg-1 | F | CTGCCTAATACCGTTGCCTTCTT | |
| | R | AATCCGGACGGCTCCAA | |
| sdhA-1 | F | TCGCAGCTCAAGGAGGAATC | 115 |
| | R | ATGGCATCCTGATCTCCGAG | |
| sdhB-1 | F | ATGCAAGCCTACAGATGGGT | 148 |
| | R | CCTTAGCTGGGTTCAAGTGTTTT | |
| sdhD-1 | F | CCAGAAGCGCTCCAAGAATC | 112 |
| | R | GCCCAAAGACGTTCAAGCTTA | |
| cdc-42 | F | GAGAAAAATGGGTGCCTGAA | 111 |
| | R | CTCGAGCATTCCTGGATCAT | |
| pmp-3 | F | GTTCCCGTGTTCATCACTCAT | 115 |
| | R | TCTACAGCTTCTCGACGGTGT | |

1. Hunter, S.E., et al., *The QPCR assay for analysis of mitochondrial DNA damage, repair, and relative copy number.* Methods, 2010. **51**(4): p. 444-51.
2. Bratic, I., et al., *Mitochondrial DNA level, but not active replicase, is essential for Caenorhabditis elegans development.* Nucleic acids research, 2009. **37**(6): p. 1817-28.
3. Rooney, J.P., et al., *PCR based determination of mitochondrial DNA copy number in multiple species.* Methods Mol Biol, 2015. **1241**: p. 23-38.
